# *Staphylococcus aureus* Mnh1 cation–proton antiporter promotes phagocytic uptake and intracellular survival in human monocytes

**DOI:** 10.64898/2026.07.29.730383

**Authors:** Aya Iizuka, Maria Vittoria Mazzuoli, Emre Demirbilek, Anne-Kathrin Woischnig, Richard Kuehl, Vincent de Bakker, Dirk Bumann, Annelies S Zinkernagel, Jan-Willem Veening, Nina Khanna

## Abstract

While *Staphylococcus aureus* is predominantly an extracellular human commensal, it can also exhibit an intracellular lifestyle that enables its persistence within diverse host cell types, including monocytes and macrophages. This capacity complicates treatment, contributing to chronic infections and therapeutic failure. Employing combined flow cytometry and microscopy, we show that monocyte internalization is associated with markedly increased antibiotic survival of intracellular *S. aureus*. To identify staphylococcal genetic determinants important for its intracellular lifestyle, we utilized a genome-wide CRISPRi-seq screening strategy using a co-infection model with human-derived THP-1 cells. This screen revealed a set of genes specifically required for intracellular fitness, confirming established virulence factors as well as identifying novel contributors to uptake and survival within host phagocytes. Among these, we identified the cation–proton antiporter Mnh1 and demonstrated that it is particularly important for phagocytic entry. Time-lapse microscopy demonstrated that its depletion significantly reduced uptake by THP-1 cells leading to reduced intracellular bacterial load. Functional assays further showed that the Mnh1 operon plays a critical role in the intracellular survival of multiple *S. aureus* lineages in THP-1 cells. Consistent with this finding, levofloxacin treatment of infected monocytes cleared knockdown strains more efficiently than the control strain. Our findings underscore the intracellular niche as a critical reservoir for bacteria and suggest that selective inhibition of Mnh1, in combination with clinically relevant antibiotics, offers a promising therapeutic approach for the treatment of intracellular *S. aureus* infections.

## Introduction

*Staphylococcus aureus* is a highly versatile and clinically significant human pathogen, responsible for diseases ranging from mild skin and soft tissue infections to severe invasive conditions such as endocarditis and sepsis, and is a leading cause of chronic and implant-associated infections^1–4^. Although antibiotic resistance contributes to its clinical impact, it does not fully account for the frequent treatment failures observed in *S. aureus* infections ^5,6^. While *S. aureus* predominantly lives as part of the commensal skin and nasopharyngeal epithelia microbiota, intracellular persistence has emerged as a central mechanism underlying chronicity and relapse of *S. aureus* disease. *S. aureus* is capable of invading and surviving within diverse host cell types, including non-professional phagocytes (e.g., epithelial, endothelial, and osteoblast cells) and professional phagocytes (e.g., macrophages and neutrophils)^7–22^. Within these intracellular niches, *S. aureus* undergoes profound physiological adaptation, enabling evasion of phagolysosomal killing and reprogramming of metabolic and transcriptional pathways in response to host-imposed stress^11,12,23–28^.

Recent in-human imaging data indicate that, in deep-seated infections characterized by recurrent treatment failures, *S. aureus* predominantly resides within host cells^29^. These intracellular bacteria localize to CD14^+^CD16⁺CD68⁺ cells, consistent with non-classical monocytes^29^, underscoring this cell population as a clinically relevant niche for persistence. Given prior evidence for strain-dependent phagocytic uptake ^30^, we selected *S. aureus* Cowan I, a clinically prevalent Clonal Complex 30 (CC30) strain with enhanced intracellular persistence, as a model to interrogate the mechanisms of intracellular survival ^27,28,31,32^.

Recent advances in high-throughput genetic screening have transformed bacterial functional genomics, particularly through transposon insertion sequencing (Tn-seq) and CRISPR interference sequencing (CRISPRi-seq)^33–39^. In contrast to Tn-seq, CRISPRi-seq employs a catalytically inactive Cas9 (dCas9) to sterically block transcription, enabling tunable, inducible gene knockdown, of all genes, including essential ones^34,35,39,40^. Moreover, the compact and designable nature of CRISPRi libraries reduces susceptibility to population bottlenecks in pooled in vitro growth and in vivo infection models, thereby improving the robustness of fitness estimates for critical targets^34^. In *S. aureus*, CRISPRi-seq has proven effective for mapping the essentialome^41,42^ and identifying condition-specific essential gene sets, such as determinants of dalbavancin susceptibility^36^ and genes required for growth in milk^43^. To date, the intracellular essentialome of *S. aureus* has primarily been investigated using Tn-seq^38,44^. While these studies validated known virulence factors and identified novel candidates, their findings have been limited by incomplete cross-strain validation and a lack of systematic functional follow-up ^38,44^. In addition, comparisons to bacteria grown in the absence of host cells have constrained the ability to resolve host-specific genetic requirements^38,44^.

Here, we overcome these limitations by directly comparing intracellular and extracellular bacterial populations during host cell co-culture, enabling a paired and physiologically relevant assessment of gene essentiality within the host environment. Using CRISPRi-seq, we identify a set of genes specifically required for intracellular survival, including multiple members of the Mnh1 operon (*mnhABCDEFG*). We further demonstrate that the Mnh1 operon contributes to bacterial internalization and supports intracellular persistence. Notably, depletion of the Mnh1 operon in combination with levofloxacin treatment results in clearance of intracellular *S. aureus*.

## Results

### Modelling *S. aureus* monocyte infection and single-cell analysis of intracellular persistence

To dissect mechanisms underpinning *S. aureus* intracellular survival in monocyte-derived cells, we established a controlled *in vitro* infection model using THP-1 monocytes. The Cowan I strain was selected based on its well-documented capacity for intracellular persistence across diverse host cell types, including epithelial cells, osteoblasts, and fibroblasts^27,28,32^. To enable quantification and visualization of intracellular *S. aureus*, we optimized an infection workflow compatible with ImageStream technology, which integrates flow cytometry with high-resolution microscopy to allow single-cell event analysis (**Figure 1a**). Intracellular *S. aureus* was detected using Vancomycin–JF669 (Van-JF669), a far-red fluorescent probe with superior signal-to-noise characteristics compared to conventional dyes such as RADA^45^. This approach enabled precise quantification of infected cells and direct visualization of intracellular bacteria across time points (**Figure 1b**). We next characterized intracellular survival dynamics by quantifying colony forming units (CFU) over a 5-hour time course in intracellular, extracellular, and monoculture conditions. While bacterial growth was observed in extracellular and monoculture conditions, intracellular CFU stabilized from 3 hours post-infection onward, consistent with bacterial persistence in the absence of replication (**Figure 1c**). To complement CFU measurements, intracellular bacterial burden was assessed by quantifying the Van-JF669-positive area per infected cell. This analysis revealed a plateau in bacterial signal after 3 hours, suggesting the establishment of a steady state in which bacterial uptake and replication are counterbalanced by host cell turnover and/or secondary infection events (**Figure 1d**). Given the prior evidence that intracellular localization protects *S. aureus* from antimicrobial activity ^9,12,19,46,47^, we next evaluated susceptibility within this niche. THP-1 cells were infected with Cowan I at a multiplicity of infection (MOI) of 10 for 1 hour, followed by treatment with daptomycin, flucloxacillin and levofloxacin. CFU were quantified from monocultures, extracellular, and intracellular compartments immediately post-infection (pre-treatment) and at 2, 4, 6, 8, and 24 hours after antibiotic addition (**Figure 1e**). As expected, all antibiotics exhibited potent bactericidal activity in monoculture. Extracellular bacteria showed a ∼3-log reduction, whereas intracellular *S. aureus* displayed markedly reduced susceptibility, failing to achieve a comparable reduction even after 24 hours of treatment (**Figure 1e**). These findings underscore the intracellular niche as a critical reservoir for bacteria and a potential driver of treatment failure. Importantly, comparable intracellular persistence phenotypes were observed across three distinct *S. aureus* lineages, including Cowan I (CC30), indicating that this model captures a broadly conserved feature of *S. aureus* biology rather than a strain-specific effect (**Figure S1**).

**Figure 1.**
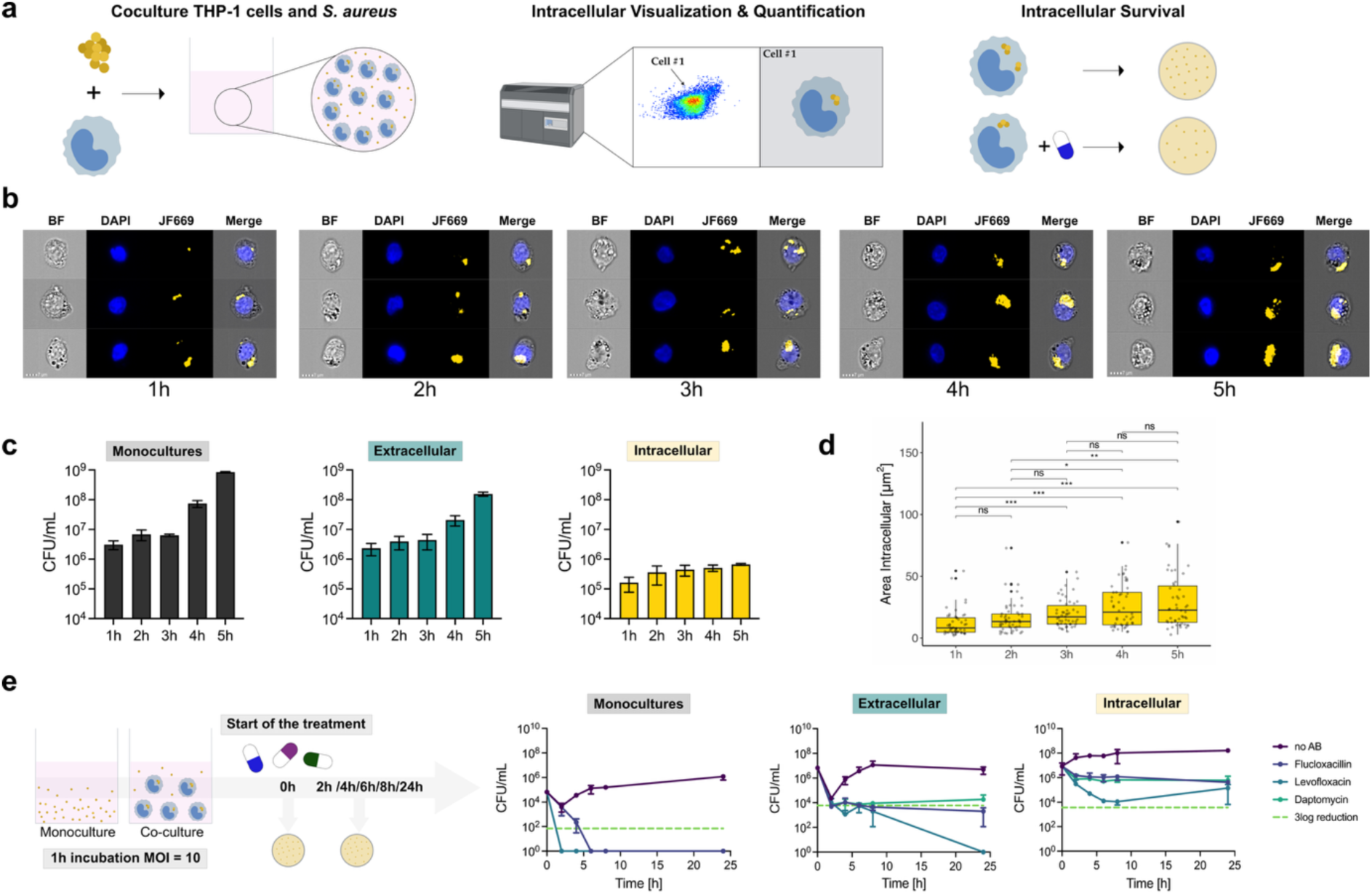
Development of a co-culture model using *Staphylococcus aureus* Cowan I and THP-1 cells. **(a)** Schematic overview of the experimental design. THP-1 cells were infected with *S. aureus* Cowan I at various time points in RPMI supplemented with 10% FBS. Intracellular bacteria were visualized and quantified using ImageStream flow cytometry, enabling single-cell event analysis. In parallel, intracellular bacterial survival under clinically relevant antibiotic exposure was assessed by colony forming unit (CFU) enumeration. **(b)** Representative images captured using the ImageStream instrument at 1 to 5 hours post-infection. Nuclei were stained with DAPI (blue), and *S. aureus* was labeled with Van-Jan669 (yellow) as described by Jantarug *et al*. (2024)^45^. At each time point, monocultures, extracellular, and intracellular bacteria were quantified, and the area of internalized *S. aureus* was measured using Fiji^48^. **(c)** CFU were quantified from co-cultures of *S. aureus* Cowan I and THP-1 cells at a multiplicity of infection (MOI) of 10, measuring both extracellular and intracellular bacterial populations, as well as monoculture controls. **(d)** Intracellular bacterial area (JF669) was quantified at multiple time points to assess phagocytic uptake dynamics. Statistical significance was assessed using the Kruskal–Wallis test followed by pairwise Wilcoxon comparisons with Holm correction for multiple testing. Asterisks indicate adjusted p-values: *** < 0.001, ** < 0.01, * < 0.05; ns, not significant. **(e)** Growth curves were generated to assess the survival of *S. aureus* over 24 hours in monoculture, and in extracellular and intracellular compartments, with or without antibiotic treatment. THP-1 cells were infected at a multiplicity of infection (MOI) of 10 and incubated for 1 hour; parallel monocultures were established using the same inoculum. Following infection, cultures were centrifuged, supernatants removed, and fresh medium—with or without antibiotics—was added. Antibiotics were applied at C_max_ concentrations, as defined by *Sanford* and/or *Mensa*^49,50^ guidelines (**Table S1**). Colony-forming units (CFU) were measured at 2, 4, 6, 8, and 24 hours post-treatment. Data shows mean of 3 biological replicates +/-SEM.

### Establishment of a genome-wide CRISPRi-seq screen in a monocyte co-infection model

To systematically identify genes required for intra- and extracellular survival, we implemented a genome-wide CRISPRi-seq screen within our co-infection model. We engineered a marker-less dCas9 variant controlled by a tetracycline-inducible promoter responsive to anhydrotetracycline (aTc) and doxycycline, integrated into the *sep* (Staphylococcal expression platform) intergenic locus of the Cowan I *S. aureus* genome. A pooled sgRNA library comprising 2,865 guides, targeting 98.6% of annotated open reading frames, was cloned into the pVL4930 vector and introduced into the Cowan I *dcas9* strain, generating a population of 2,865 knockdown mutants^42^ (**Figure 2a**). To confirm functionality of the CRISPRi system under infection-relevant conditions, we assessed gene repression in RPMI supplemented with 10% FBS in comparison with TSB. A strain carrying an sgRNA targeting the essential gene *pbpA*, encoding penicillin-binding protein 1 (PBP1)^51^, exhibited marked growth impairment upon induction, as measured by optical density measurements (OD_660_) (**Figure S2**). Consistently, genome-wide CRISPRi-seq analysis in this medium defined the baseline gene requirements for bacterial growth (**Figure S3**).

**Figure 2.**
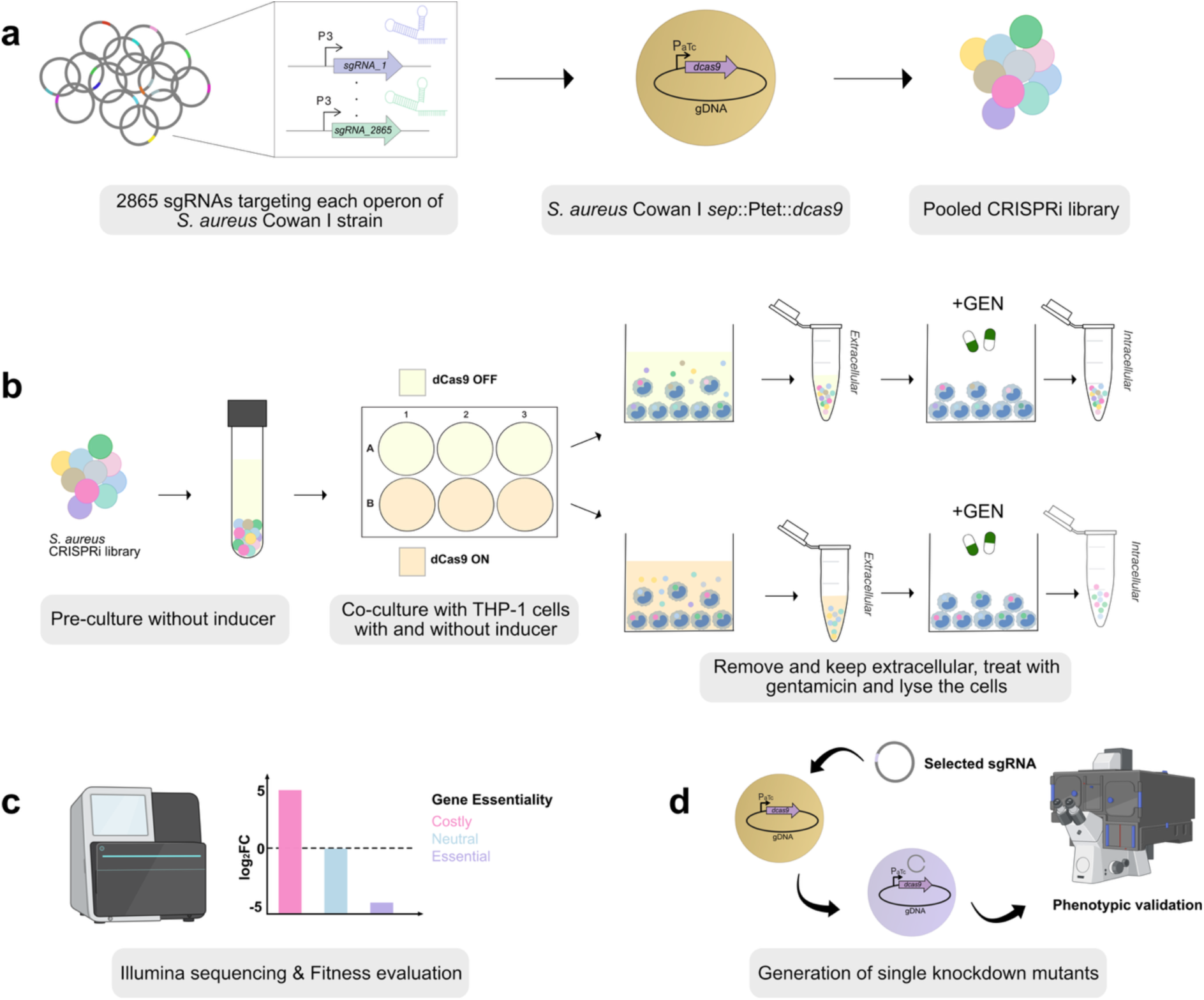
Experimental workflow overview. **(a)** A pooled CRISPR interference (CRISPRi) library was constructed to target 2,865 genes in the *S. aureus* Cowan I^42^. The *dcas9* gene was chromosomally integrated at the intergenic locus *sep* and placed under the control of a Ptet-inducible promoter. **(b)** The library was pre-cultured without the inducer (dCas9 OFF) to allow uniform growth of knockdowns. THP-1 cells were then infected with the pre-culture at a multiplicity of infection (MOI) of 10. Co-cultures were either left uninduced or treated with anhydrotetracycline (aTc) to induce *dcas9* expression (dCas9 ON). After 5 hours of infection, extracellular bacteria were harvested, and intracellular bacteria were isolated following a 1-hour treatment with gentamicin (50 μ g/mL) to eliminate residual extracellular bacteria. THP-1 cells were subsequently lysed to recover intracellular bacteria. **(c)** Plasmids encoding sgRNAs were extracted from bacterial populations and sequenced. Fitness effects were analyzed using DESeq2^53^ **(d)** sgRNAs required for intracellular survival were identified, and corresponding single-knockdown mutants were generated to validate intracellular fitness using confocal microscopy. All conditions were performed in quadruplicates. Biorender.com was used to generate part of the figure.

We next applied the CRISPRi library to the THP-1 co-infection model. Cells were infected with a multiplicity of infection of 10 (MOI=10) for 5 h in the presence or absence of inducer (**Figure 2b**). This approach enabled the selection of genes contributing not only to intracellular survival but also to host cell entry, as phagocytosis proceeds dynamically throughout the co-culture period. Because maintenance of the CRISPRi system requires chloramphenicol (10 µg/mL) for sgRNA plasmid selection and aTc (10 ng/mL) for dCas9 induction, we evaluated their impact on host cell viability and apoptosis. THP-1 cells exposed to concentrations 10-fold higher than those used in the assay showed no increase in cell death or apoptosis compared to untreated controls, confirming the compatibility with the co-culture system (**Figure S4**).

Following infection, extracellular bacteria were collected, while THP-1 cells were washed and treated with gentamicin to ensure the intracellular population was not contaminated by extracellular bacteria^11,15,52^. (**Figure 2b**). Cells were subsequently lysed to recover intracellular bacteria. Plasmid DNA was extracted and sgRNA abundance was quantified by sequencing. Differential enrichment analysis was then performed to infer gene-specific fitness contributions under intracellular and extracellular conditions (**Figure 2c**). To validate the screen, specific hits were chosen for the generation of single knockdown mutants (**Figure 2d**).

### CRISPRi-seq identifies genes required for intracellular and extracellular survival

sgRNA count profiles were consistent within growth condition, and aTc addition both explained most of the variance between samples and produced a pronounced depletion of a subset of sgRNAs, consistent with efficient CRISPRi induction (**Figure 3a, Figure S5-6**). Of note, extracellular samples showed greater variability at the sgRNA count totals and a relative enrichment of sgRNAs in uninduced normalized counts (**Figure S5c**). This might be due to slight sampling bias during supernatant (extracellular bacteria) pipetting, rather than a systematic effect. This is supported by the white gaps observed in the heatmap (**Figure S6**) which reflects random, replicate-specific depletion of sgRNA subsets. Despite this background activity, clear differences in sgRNA enrichment between uninduced and induced samples were observed within each condition (**Figure S6**).

**Figure 3.**
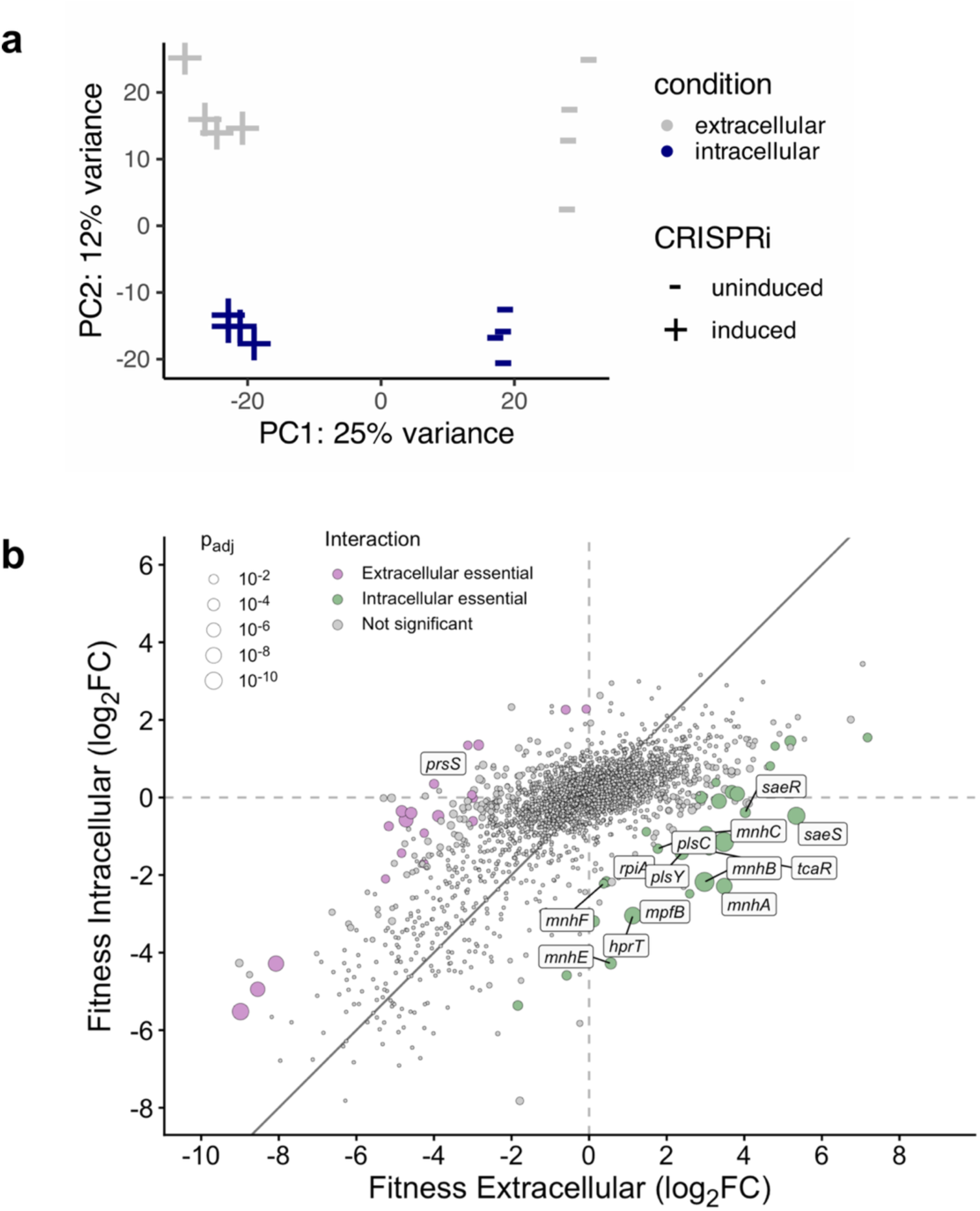
Comparative analysis of intracellular and extracellular essentialomes. **(a)** Principal Component Analysis (PCA) highlighting the separation between intracellular and extracellular samples. The first principal component (PC1), accounting for 25% of the variance, reflects the CRISPRi induction status (+aTc vs. –aTc), while the second component (PC2), explaining 12% of the variance, reflects the compartment-specific condition (intracellular vs. extracellular). **(b)** Interaction plot showing the differential impact of CRISPRi induction between intracellular and extracellular conditions. Genes associated with fitness loss when repressed in the extracellular environment are shown in pink, while those associated to fitness loss intracellularly are shown in green. FDR-adjusted p-values testing for a difference in log_2_FC between the two conditions are shown.

Principal Component Analysis (PCA) demonstrated a clear separation of induced intracellular and extracellular samples, indicating distinct condition-specific genetic requirements (**Figure 3a**). This divergence aligns with prior observations that intracellular *S. aureus* adopts a metabolically and physiologically distinct state compared to its extracellular counterpart, including altered ATP levels and adaptive metabolic reprogramming during persistence^12,54,55^.

To define genes with differential fitness across infection environments, we compared fitness scores by calculating the difference in log₂ fold change (log₂FC) between intracellular and extracellular conditions (log₂FC_intracellular_ − log₂FC_extracellular_). This analysis revealed a distinct subset of genes with significant differential fitness in either niche, demonstrating that CRISPRi-seq robustly resolves condition-specific essentialomes (**Figure 3b**, **Table 1**). Among the genes preferentially required for intracellular survival, we identified the *saeRS* two-component global regulatory system. *saeRS* has been implicated in host-pathogen interactions, including epithelial cell adhesion, internalization, and survival within neutrophils^56,57^. Consistent with prior studies, loss of *saeRS* function is associated with increased susceptibility to macrophage-mediated killing and impaired phagosomal escape, supporting its central role in intracellular adaptation ^25,26^. We also identified *prsS* as specifically required for extracellular survival. This gene encodes a membrane-associated stress response protein previously linked to virulence in both human and murine infection models, highlighting distinct stress adaptation requirements between extracellular and intracellular environments ^58,59^. Together, these findings validate the capacity of our screening approach to resolve niche-specific genetic requirements and uncover previously uncharacterized determinants of *S. aureus* intracellular survival and host cell internalization.

**Table 1.**
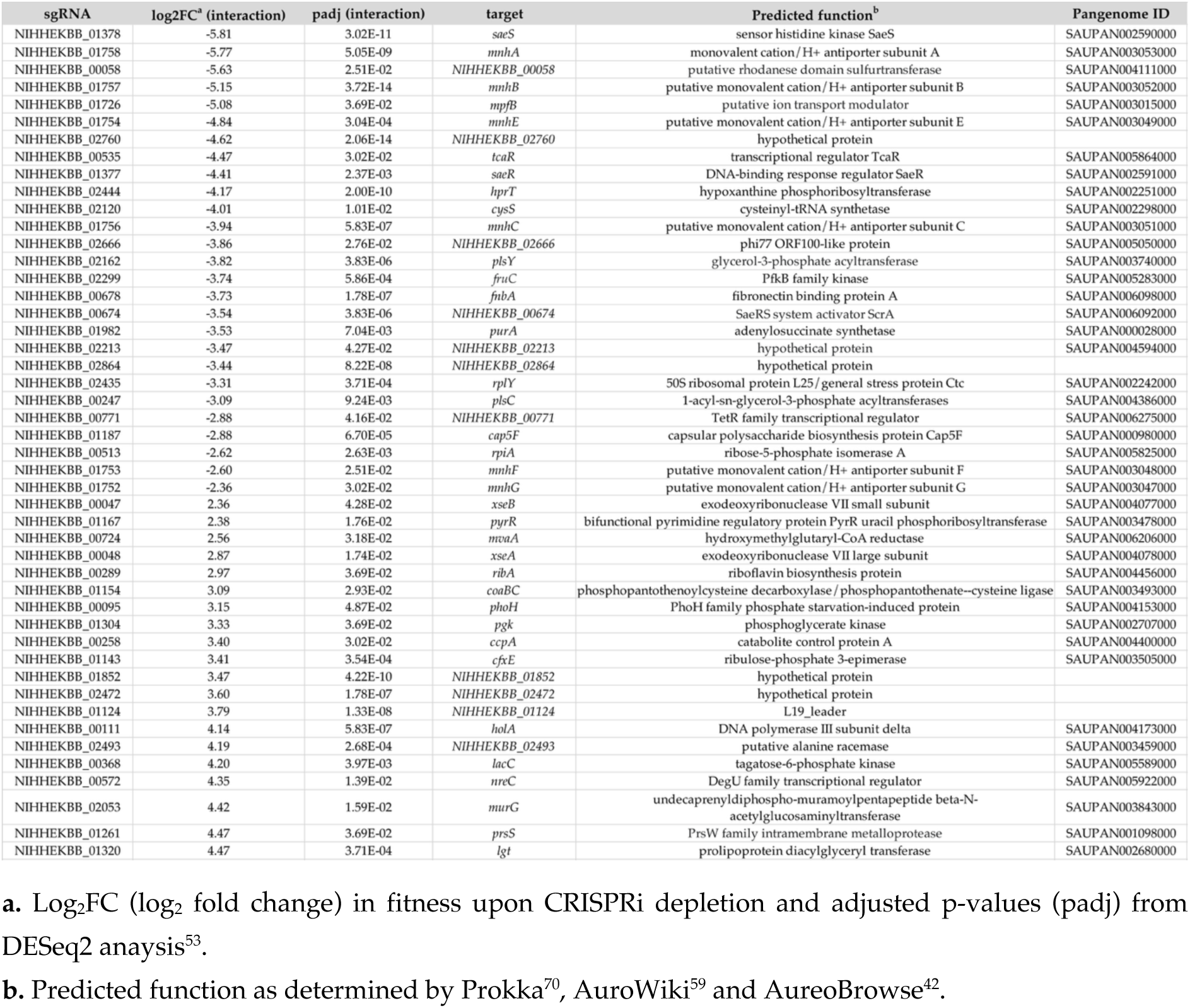
Genes with differential fitness effects in intracellular or extracellular environments as determined by CRISPRi-seq.

### Genes specifically required for the intracellular lifestyle of *S. aureus*

Focusing on intracellular persistence, we next interrogated genes defined by reduced fitness under intracellular conditions but non-deleterious effects extracellularly. This analysis identified a set of candidate genes selectively required for survival within host cells, providing a foundation for targeted functional validation (**Figure 3b**, **Table1, Table S2**). Notably, genes involved in phospholipid biosynthesis, including *plsY* and *plsC*, were among the top intracellular-specific hits. These enzymes catalyze early steps in the phosphatidic acid synthesis, a key precursor for membrane phospholipids. A nonsynonymous single nucleotide polymorphism (SNP) in *plsY* has been associated with reduced growth rate and attenuated virulence in *S. aureus*^60^. While direct evidence of *plsC* essentiality in *S. aureus* is lacking, this gene has been shown to be essential in other bacterial species, including *E. coli*^61^.

The transcriptional regulator *tcaR* was also identified as an intracellular fitness determinant. TcaR modulates expression of cell surface and biofilm-associated genes, including *spa*, which encodes protein A^62–64^. Given the role of protein A in immune evasion through binding to the Fc region of immunoglobulins, altered regulation of cell surface architecture may influence bacterial uptake, immune recognition, or intracellular survival.

Genes involved in nucleotide metabolism, including *hprT* and *rpiA*, were also selectively required intracellularly. Although not previously linked to *S. aureus* intracellular survival, studies in other pathogens suggest that intracellular environments are depleted in nucleotide precursors, necessitating efficient salvage and biosynthesis pathways. The requirement for *hprT*, which mediates purine salvage, may reflect an increased demand for nucleotide recycling under conditions of DNA damage and replication stress, while *rpiA* supports nucleotide biosynthesis through the pentose phosphate pathway ^65^ ^66,67^ ^68^. Another gene that was significantly only essential for intracellular survival was *mpfB*, a protein involved in magnesium export. It is regulated by Sigma B and was shown to compensate for *mpfA* under stress condition^69^. Chang et al (2025)^38^ recently showed that *mpfA* was specifically essential for intracellular *S. aureus* survival in non-professional phagocytic compared to blood and *in vivo* condition. It supports the importance of magnesium metabolism in intracellular survival. Importantly, the Mnh1 operon (*mnhA– mnhG*) encoding a multi-subunit cation/proton antiporter, emerged as a major determinant of intracellular survival, with multiple components showing strong fitness defects. This finding is supported by recent independent studies demonstrating its importance in post-entry survival in epithelial cells, underscoring a conserved role in intracellular adaptation ^38^.

Based on these results, we prioritized the Mnh1 operon and generated an *mnhB* knockdown strain for further characterization. Notably, until recently, this operon had not been directly linked to intracellular survival in monocytes, highlighting the power of our approach to uncover previously unrecognized determinants of host–pathogen interactions.

### *mnhB* promotes early intracellular entry across diverse *S. aureus* lineages

To validate the role of *mnhB*, we generated knockdown strains in three distinct *S. aureus* lineages. Growth in cell culture medium with or without aTc revealed no marked fitness defect in the absence of THP-1 cells for any lineage (**Figure 4a-c**) indicating that *mnhB* is dispensable under these conditions.

**Figure 4.**
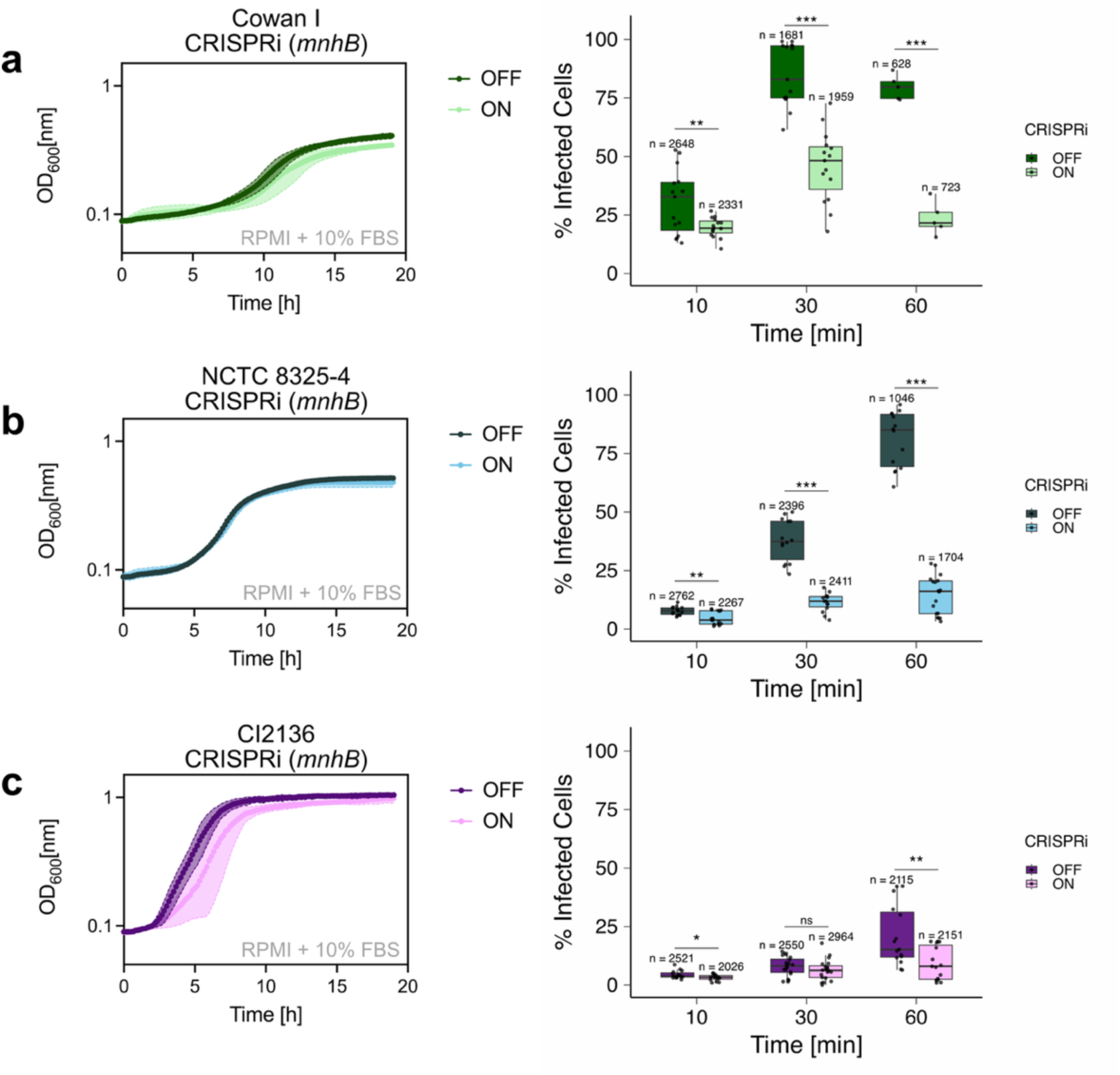
*mnhB* contributes to early intracellular entry of *S. aureus* into THP-1 cells. **a–c**, Knockdown of *mnhB* using an inducible CRISPRi system (ON: +aTc, 10 ng/mL; OFF: –aTc) was performed in three *S. aureus* strain backgrounds: **(a)** Cowan I, **(b)** NCTC 8325-4, and **(c)** clinical isolate CI2136. Prior to infection, bacterial cultures were grown for 5 hours with or without induction in RPMI + 10% FBS to assess potential growth defects. THP-1 cells were infected at a multiplicity of infection (MOI) of 10 and incubated for 10, 30, or 60 minutes. After infection, extracellular bacteria were removed by washing, and cells were fixed, stained, and imaged by confocal microscopy to compare intracellular bacterial abundance between uninduced and induced knockdown conditions. THP-1 cytoplasm was stained with CellTracker to facilitate accurate quantification of intracellular *S. aureus* by Z-stack confocal microscopy (see **Methods**). Each data point represents the mean percentage of infected cells per field of view, each fields containing ≥70 cells. For each condition, 3–5 images were acquired per experiment, and experiments were independently repeated 3–4 times. Statistical significance was determined using the Wilcoxon rank-sum test. Asterisks denote p-values: *** < 0.001, ** < 0.01, * < 0.05; ns, not significant. The total number of cells quantified per condition is shown above each boxplot.

We next assessed the contribution *mnhB* to host cell entry using short-term co-culture infections with pre-induced mutants. THP-1 cells were infected for 10, 30, and 60 minutes, and the proportion of infected cells was quantified by confocal microscopy. Baseline phagocytosis rates differed across lineages, with higher uptake observed in the Cowan I strain compared to NCTC 8325-4 and CI2136. Despite this variability, *mnhB* depletion consistently reduced infection levels across all backgrounds and time points (**Figure 4**). In both Cowan I and NCTC 8325-4 backgrounds, *mnhB* depletion resulted in a significant reduction in infected cells at all time points (**Figure 4a–b**). In the clinical isolate CI2136, a similar decrease was observed at 10 and 60 minutes, but not at 30 minutes (**Figure 4c**). Control experiments conducted in strains lacking the sgRNA plasmid showed no differences in the number of infected cells, confirming that the phenotype is specific to *mnhB* depletion (**Figure S7**). Together, these results identify *mnhB* as a conserved determinant of early intracellular entry in *S. aureus*, independent of strain-specific differences in uptake efficiency.

### *mnhB* knockdown reduces intracellular bacterial burden across lineages

To complement infection rate measurements, we quantified intracellular bacterial burden by assessing the JF669 fluorescence signal per cell at the same time points. Across all three lineages, *mnhB* depletion led to a significant reduction in mean intracellular bacterial volume at every time point compared to uninduced controls (**Figure 5a–c**, **Figure 6a-b**). These findings indicate that, beyond impairing host cell entry, *mnhB* knockdown may also compromises intracellular persistence of *S. aureus*. Control experiments conducted in strains lacking the sgRNA plasmid showed no differences in intracellular bacterial volume, confirming that the observed phenotype is specific to *mnhB* depletion **(Figure S8).**

**Figure 5.**
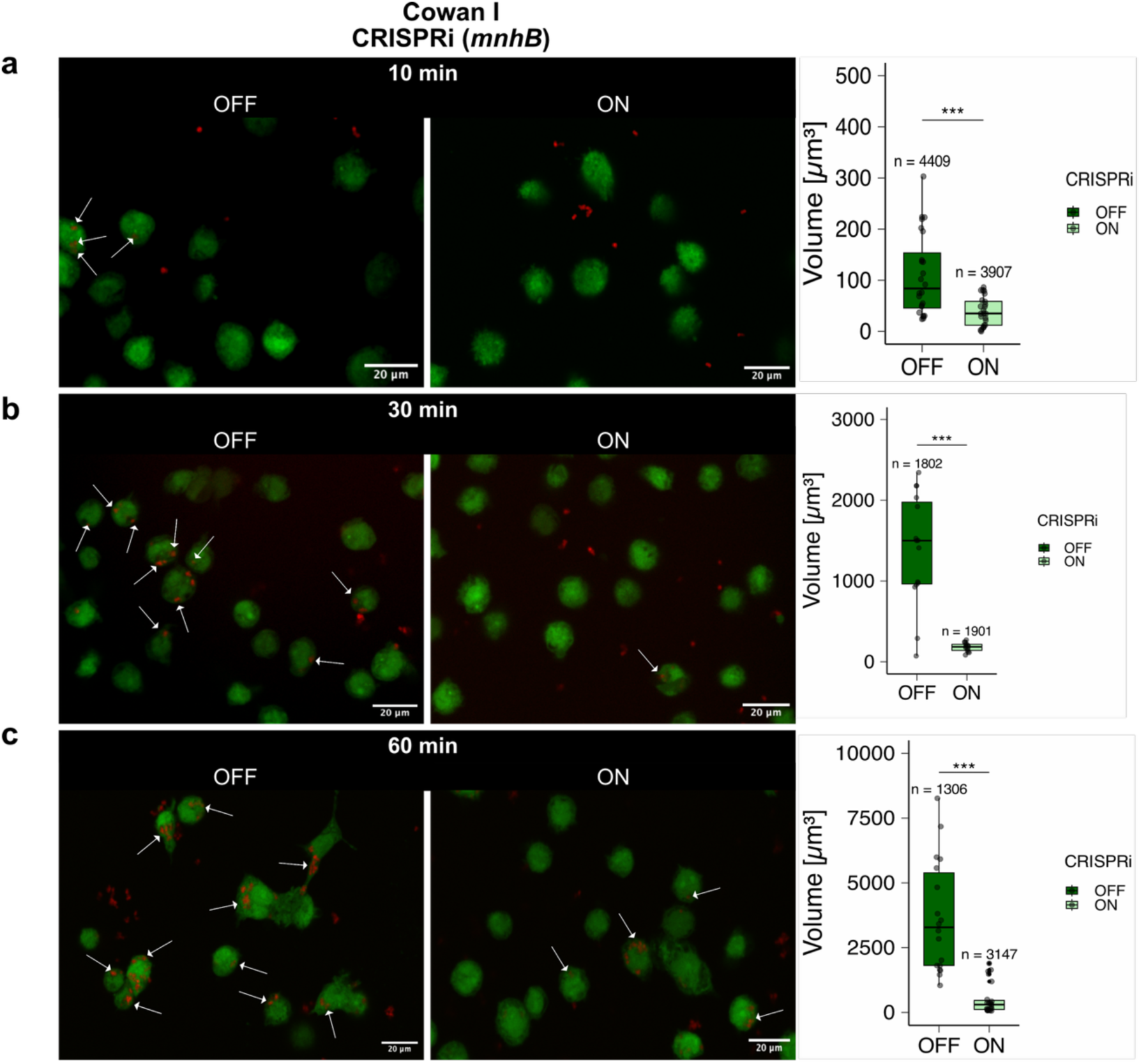
Depletion of *mnhB* reduces the intracellular volume of *S. aureus* Cowan I in THP-1 cells. **(a)** 10 min, **(b)** 30 min, and **(c)** 60 min. In addition to quantifying the proportion of infected cells (Figure 4), we measured the intracellular bacterial volume per cell. For each time point, representative images with and without aTc are shown. White arrows indicate cells containing intracellular *S. aureus*. Each data point represents the mean percentage of bacterial volume per cell within a field of view, with each field containing ≥70 cells. For each condition, 3–5 images were acquired per experiment, and experiments were independently repeated 3–4 times. Statistical significance was assessed using the Wilcoxon rank-sum test. Asterisks indicate *p*-values: *** < 0.001, ** < 0.01, * < 0.05; ns, not significant. The total number of cells quantified per condition is shown above each boxplot.

**Figure 6.**
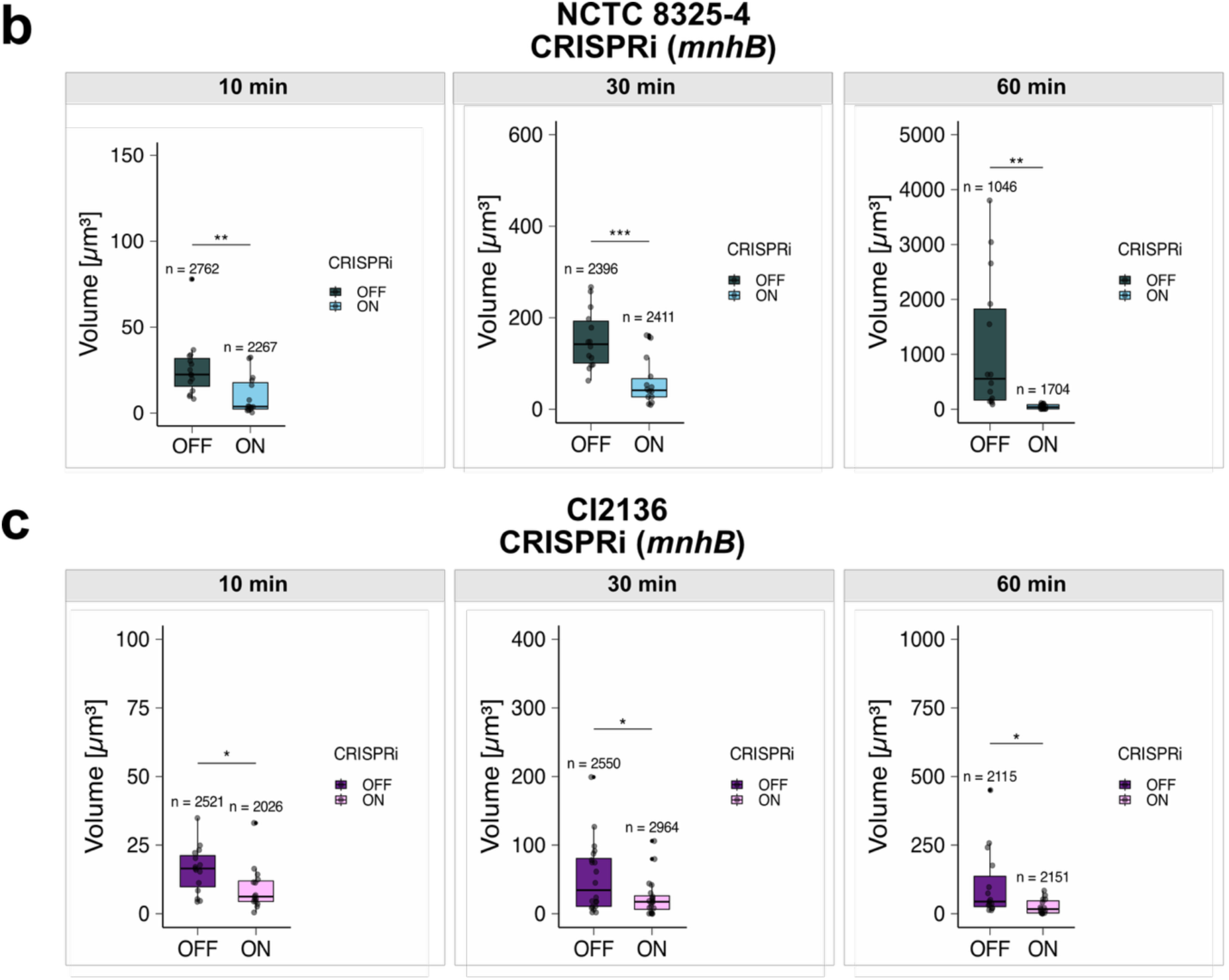
Depletion of *mnhB* reduces the intracellular volume of *S. aureus* NCTC 8325-5 and CI2136. **(a)** Intracellular bacterial volume measured at 10, 30, and 60 min in NCTC 8325-5. **(b)** Intracellular bacterial volume measured at 10, 30, and 60 min in clinical isolate CI2136. Each data point represents the mean percentage of bacterial volume per cell within a field of view, with each field containing ≥70 cells. For each condition, 3–5 images were acquired per experiment, and experiments were independently repeated 3–4 times. Statistical significance was assessed using the Wilcoxon rank-sum test. Asterisks indicate *p*-values: *** < 0.001, ** < 0.01, * < 0.05; ns, not significant. The total number of cells quantified per condition is shown above each boxplot.

### Combined *mnhB* depletion and levofloxacin treatment enables complete clearance of intracellular *S. aureus*

Given the reduced intracellular burden observed upon *mnhB* knockdown, we tested whether targeting the Mnh1 operon could enhance antibiotic efficacy. We therefore combined *mnhB* depletion with levofloxacin treatment and monitored intracellular bacterial survival (**Figure 7a**). Importantly, this combination resulted in complete clearance of intracellular *S. aureus* in induced knockdown strains, whereas uninduced controls retained persistent bacterial populations. This effect was consistently observed across all three lineages tested (**Figure 7b–d**), indicating that disruption of Mnh1 function sensitizes intracellular bacteria to antibiotic-mediated killing.

**Figure 7.**
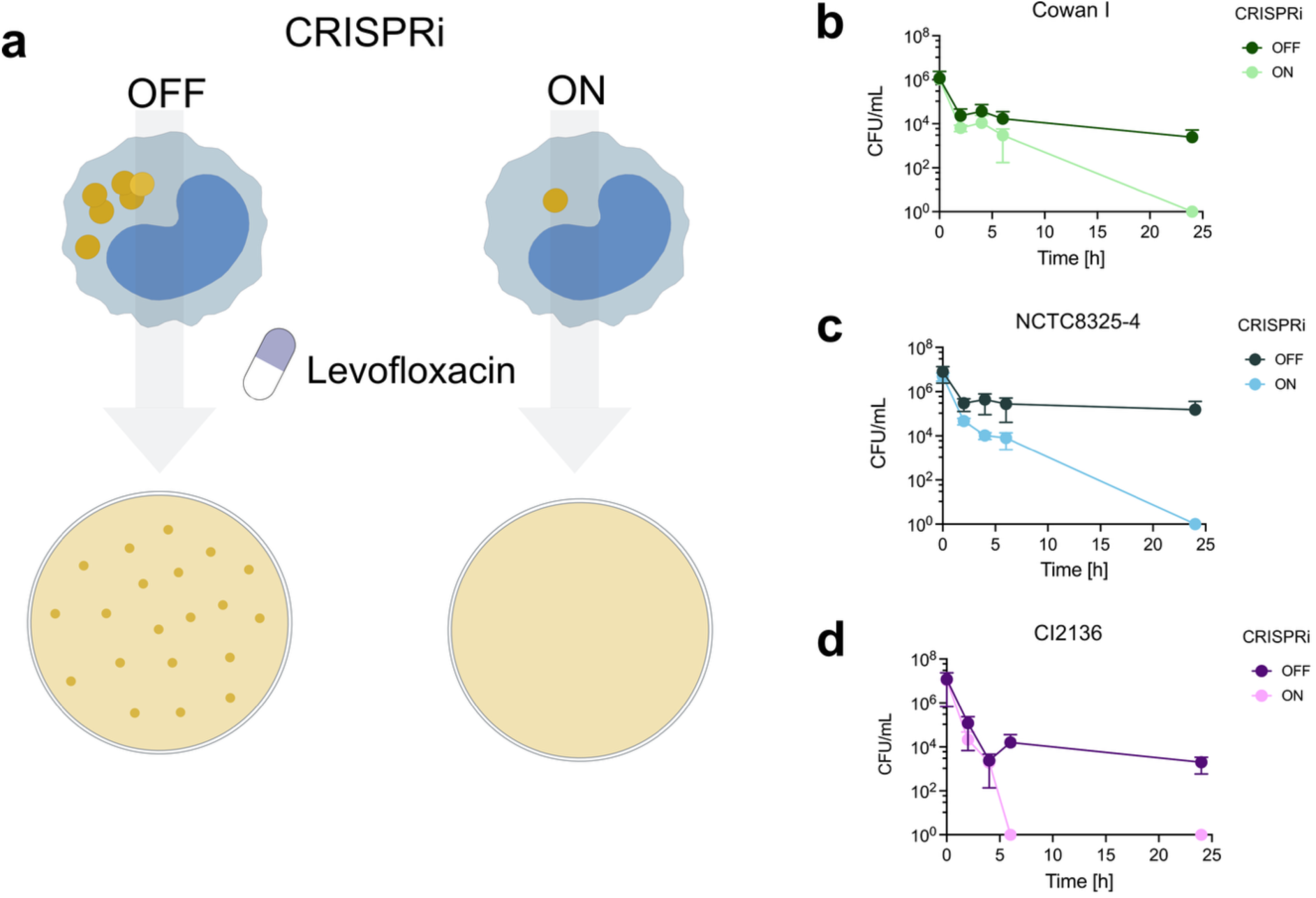
Levofloxacin treatment of infected THP-1 cells with *mnhB* depletion enhances intracellular bacterial clearance in three different *S. aureus* lineages. Time-kill assays with levofloxacin were performed on the *S. aureus* strain carrying the sgRNA-*mnhB*. **(a)** Intracellular CFU counts were measured for the knockdown strains under induced and uninduced conditions following levofloxacin exposure. **(b)** Killing curves comparing OFF and ON conditions in **(b)** Cowan I (ST30), **(c)** NCTC8325-4 (ST8), **(d)** CI2136 (ST45).

## Discussion

*S. aureus* remains a leading cause of morbidity and mortality worldwide, in part due to its capacity to persist within host cells and evade antibiotic treatment. Here, using a genome-wide CRISPRi-seq approach in a physiologically relevant co-infection model, we identify the Mnh1 Na⁺/H⁺ antiporter operon as a key determinant of the intracellular lifestyle of *S. aureus*.

Functional validation using an ImageStream-based analysis demonstrates that depletion of *mnhB*, and consequent disruption of the operon, impairs both early host cell entry and intracellular persistence across multiple strain backgrounds. These findings position Mnh1 as a dual-function factor linking initial host–pathogen interactions with adaptation to the intracellular niche. Mechanistically, the observed phenotypes likely reflect disrupted ion homeostasis. Na⁺/H⁺ exchangers are central to maintaining cytoplasmic pH, membrane potential, and ionic balance ^71,72^. Perturbation of these processes may alter cell surface properties and cell wall architecture ^73,74^, thereby affecting adhesin function and reducing bacterial uptake. We hypothesize that once internalized, loss of Mnh1 function may compromise the ability of *S. aureus* to counteract phagosomal acidification and associated stresses, leading to impaired metabolic adaptation and reduced intracellular survival. These findings are consistent with previous work highlighting the importance of ion transport systems for intracellular bacterial fitness^38^.

Importantly, targeting Mnh1 also sensitized intracellular *S. aureus* to antibiotic treatment. Combination of *mnhB* depletion with levofloxacin therapy resulted in complete clearance of intracellular bacteria across all tested lineages, whereas control populations persisted. This enhanced susceptibility likely arises from the convergence of antibiotic stress with impaired homeostatic control. Disruption of Na⁺/H⁺ exchange is expected to destabilize the proton motive force, reduce ATP production, and compromise energy-dependent processes, including DNA repair and stress adaptation^75–77^. In addition, fluoroquinolone resistance in *S. aureus* is partly mediated by proton motive force–dependent efflux pumps such as NorA, NorB, and MepA^78–80^. Impairment of these systems would further increase intracellular antibiotic accumulation and bactericidal activity. Consistent with this model, disruption of membrane potential is known to reduce efflux efficiency and create synergistic vulnerabilities when combined with antibiotic treatment^81–83^.

Together, our findings identify Mnh1 as a critical vulnerability in intracellular *S. aureus* and suggest that targeting ion homeostasis can potentiate antibiotic efficacy. This concept aligns with broader efforts to overcome intracellular persistence through combination strategies. Approaches such as antibiotic–adjuvant pairing, host-directed therapies, antibody–antibiotic conjugates, cell-penetrating enzybiotics^84^ and nanoparticle-based delivery systems have shown promise in preclinical models^85–88^, although their long-term safety and comparative effectiveness remain insufficiently defined. For example, Yang et al. (2018)^85^ demonstrated enhanced intracellular killing using gentamicin-loaded nanoparticles; similar strategies could be adapted to deliver molecules that specifically interfere with *mnhB* expression or Mnh1 activity. Clinical development of nanoparticle-based delivery platforms is ongoing and represents a promising avenue to target determinants of intracellular persistence^89^.

In summary, this study establishes genome-wide CRISPRi-seq in a host–pathogen infection model as a powerful platform to resolve niche-specific bacterial vulnerabilities and identifies the Mnh1 antiporter as a key mediator of intracellular persistence in different genetic backgrounds including a clinical isolate. Targeting this system may provide a strategy to overcome treatment failure in persistent *S. aureus* infections.

## Methods

### Bacterial strains

*S. aureus* isolates used in this study included Cowan I (ATCC 12598), NCTC-8325-4, and CI2136 (clinical isolate from the University Hospital of Zürich). The clinical isolate was obtained from a case of bacteremia (ethics approval BASEC 2017-02225). Bacterial strains were revived from glycerol stocks by streaking onto agar plates and subsequently transferred to RPMI supplemented with 10% FBS (Thermo Fisher Scientific) for overnight culture. CRISPRi libraries were initiated by inoculating 50 µL of glycerol stock into 50 mL of fresh THP-1 medium supplemented with 50 µg/mL chloramphenicol, and incubated at 37 °C with shaking at 150 rpm.

### Cytotoxicity Assay

THP-1 cells were incubated in RPMI supplemented with 10% FBS and varying concentrations of chloramphenicol and/or aTc for 6 h at 37 °C and 5% CO₂. Cells were centrifuged at 360 × *g*, washed with PBS, and stained for the apoptosis marker caspase-3/7 using CellEvent™ Caspase-3/7 Green Detection Reagent (ThermoFisher) in PBS at 37 °C for 45 min. Following incubation, cells were washed and stained for viability using Zombie Aqua™ Fixable Viability Kit (ThermoFisher) in PBS at room temperature for 20 min. Cells were then centrifuged and resuspended in fixing buffer containing 4% paraformaldehyde for 20 min at room temperature. Samples were acquired on a CytoFLEX flow cytometer (Beckman Coulter), and data were analyzed using FlowJo software (BD Biosciences).

### Cloning of *dcas9* into *S. aureus* Cowan I chromosome

A markerless dCas9 variant was inserted into a newly designated intergenic *sep* locus in the *S. aureus* Cowan I genome. The *sep* locus is a conserved intergenic region flanked by *SAUPAN006480000 and SAUPAN006481000*.

### CRISPRi-seq screening

#### Library pre-cultures

The dCas9 Cowan I library was cultured overnight at 37 °C with agitation. Pre-cultures were prepared by diluting overnight cultures to an OD_600_ of 0.01 in 50 mL of fresh THP-1 medium and grown at 37 °C with 5% CO₂ for approximately 3 h, until reaching OD_600_ ≈ 0.3. Cultures were maintained in RPMI + 10% FBS + 50 µg/mL chloramphenicol without shaking, to mimic the conditions of subsequent infection assays in which shaking is not possible due to the presence of THP-1 cells. No inducer was added during this step to allow homogeneous growth of the pooled library.

#### Induction of libraries and infection

For control conditions, bacteria were grown in THP-1 medium in the presence or absence of inducer. For infection assays, THP-1 cells were co-cultured with bacteria at a multiplicity of infection (MOI) of 10 for 5 h, with or without induction by anhydrotetracycline (aTc). Extracellular bacteria were removed and cells were treated with gentamycin 50 µg/mL for 45 min at 37°C and 5% CO_2_.

### Library construction & Sequencing

Samples from screening cultures were centrifuged into pellets (4000 x g, 10 min, 4°C) and stored in TSB with 20% glycerol at -80°C. For bacterial lysis, the frozen cells were resuspended in 500 μL of Tris-EDTA buffer. 0.8 mg/mL of lysozyme and 0.02 mg/mL of lysostaphin were added and the samples were incubated for 1h at 37°C. For plasmid-isolation, a miniprep kit for *S. aureus* (PureYield Plasmid Miniprepkit System, Promega) was used. DNA was stored at -20°C. sgRNAs from the minipreps were amplified using Nextera DNA Index primers, each containing an Illumina barcode. Different primer pairs were used for the various samples from different growth conditions, allowing accurate assignment of sequencing reads to the respective condition. The amplification of sgRNAs was performed through 25 cycles of PCR. The PCR-products were purified using an agarose gel extraction kit (PCR clean-up Gel extraction, Macherey-Nagel) and quantified by Qubit DNA quantification. The samples were then normalized in RSB, pooled and sent to the UNIL Genomic Technologies Facility for sequencing. The pooled samples were sequenced on the NovaSeq platform using 150 bp reads.

### Read quality assessment, sgRNA counts and data analysis

Read quality was assessed using the 2FAST2Q (v. 2.8.1) software ^34,90^. sgRNA spacer occurrences were counted using 2FAST2Q, allowing for one mismatch and ensuring a minimum Phred score of 30 throughout the 20-nt spacer sequence. The count data of sgRNAs were analyzed with the DESeq2 package (v. 1.50.2) in R (v. 4.5.3)^53^. We defined thresholds of ± 1 for log_2_FC (testing for a significant difference from halving or doubling of an sgRNA in induced versus non-induced samples) and 0.05 for the FDR-corrected, adjusted p-values. Log_2_FC values were shrunk either with the apeglm or ashr method^91^.

### Flow Cytometry Imaging (ImageStream)

Imaging-based flow cytometry was performed using the Amnis ImageStream® X Mk II system, equipped with three objectives (20×, 40×, and 60×), 12 detection channels, and four excitation lasers (405, 488, 561, and 642 nm). Data acquisition and analysis were conducted using IDEA® (Image Data Exploration and Analysis Software), applying gating strategies to select single, in-focus cells. THP-1 monocytes were counted, and 2×10^5^ cells were seeded into each well of a 96-well plate. Cells were infected at a multiplicity of infection (MOI) of 10 and incubated at 37 °C with 5% CO₂ for the indicated time points. Following incubation, cells were washed with fresh mPBS (ThermoFisher Scientific) and fixed for 20 min at RT. Permeabilization and blocking of nonspecific staining were performed by incubating the cells in PBS-T supplemented with 10% goat serum (ab741, Abcam) for 1 h in the dark. Within the blocking solution, 0.5% Vancomycin-JF669^45^ and 0.5% DAPI (ab228549, Abcam) were added, and cells were stained for 20 min in the dark. Samples were then washed twice with PBS and imaged using the Cytek Amnis ImageStream® Mk II. Vancomycin-JF669 and DAPI were excited with 150 mW and 50 mW lasers, respectively (Jantarug et al., 2024). Events were gated for single cells based on size and aspect ratio, followed by selection of DAPI-positive and Vancomycin-JF669–positive populations to obtain images and quantitative data of cells containing intracellular *S. aureus*.

### sgRNA cloning for knockdown mutant construction

To clone sgRNAs into the pVL4930 vector via primer annealing, 17 μL of plasmid DNA was digested with 1 μL of BsmBI (NEB) and 2 μL of 10× LDR2 buffer in a total volume of 20 μL. The digestion was performed at 37 °C for 2 hours. The products were separated on a 1% agarose gel in 1× TBE buffer and visualized using SYBR Safe stain (Thermo Fisher Scientific). The linearized backbone, corresponding to the removal of the mCherry cassette, was excised and purified. The DNA was eluted in 30 μL of molecular biology-grade water (ddH₂O). Oligonucleotides corresponding to each sgRNA were designed and annealed by mixing 2.5 μL of forward and reverse primers (100 μM each) with 5 μL of 10× TEN buffer and 40 μL of ddH₂O. The mixture was incubated at 95 °C for 5 minutes and then allowed to cool gradually to room temperature. Ligation reactions were assembled by combining 2 μL of digested plasmid, 2 μL of annealed oligonucleotides (previously diluted 1:100 in ddH₂O), 4 μL of ddH₂O, 1 μL of T4 DNA ligase, and 1 μL of ligase buffer. Reactions were incubated overnight at 16 °C.

### Transformation in *E. coli*

The plasmid construct was first introduced into *E. coli* IM08B for amplification via standard bacterial cloning. Transformation was carried out by adding 10 μL of the ligation reaction to 100 μL of chemically competent *E. coli* IM08B cells. The mixture was incubated on ice for 30 minutes, followed by a heat shock at 37 °C in a water bath for 4 minutes. Subsequently, 900 μL of LB medium was added, and the cells were incubated at 37 °C with shaking for 1 hour and 30 minutes to allow recovery. The transformed cells were then plated onto LB agar plates containing the appropriate antibiotic (100 μg/mL Ampicillin) and incubated overnight at 37 °C. Successful transformation was assessed by colony PCR using Taq Plus Master Mix II (Vazyme, cat# P112-01) with primers OVL5725 and OVL7126 (**Table S3**). Only white colonies were tested, as cloning of sgRNA replace mCherry sequence in pVL4930^42^. PCR reactions were performed on a ProFlex PCR System using the following cycling conditions: initial denaturation (94°C, 5 min.), denaturation (94°C, 20 sec.), annealing (55°C, 30 sec.), extension (72°C, 30sec.), final extension (72°C, 10 min.). Denaturation, annealing and extension were repeated for 30 cycles. Amplification of the DNA was verified by agarose gel electrophoresis. PCR products were loaded onto a 1% agarose gel prepared in 1× TBE buffer and run at 120 V for 15 minutes. DNA bands were visualized using SYBR Safe stain, with the expected product size of 480 bp. Following verification, PCR products were purified using NucleoSpin silica mini-columns (Macherey-Nagel). Two volumes of NT1 binding buffer were added to each PCR reaction, and the mixture was transferred to the columns and processed under vacuum. Columns were washed twice with 650 μL of NT3 wash buffer, also under vacuum. To dry the silica membrane, columns were centrifuged at 11,000 rpm for 2 minutes. DNA was eluted with 15 μL of molecular biology grade water and recovered by a final centrifugation at 11,000 rpm for 1 minute. Purified DNA was submitted for Sanger sequencing using primer OVL867 (Table S3).

### Transformation in *S. aureus*

Plasmids carrying the sgRNA constructs were isolated from successfully transformed *E. coli* clones using the PureYield Plasmid Miniprep System (Promega, cat# A1223), according to the manufacturer’s protocol. Purified plasmid DNA was stored at −20 °C until further use. For transformation into *Staphylococcus aureus* strains Cowan I harboring dCas9, 4 μg of plasmid DNA were mixed with 100 μL of chemically competent *S. aureus* cells. Electroporation was performed in 2 mm cuvettes using the ECM 399 Electroporation System (BTX Harvard Apparatus) at 2.1 kV, 100 Ω, and 25 μF. Immediately following the pulse, 900 μL of TSB medium supplemented with sorbitol (500 mM) was added to stabilize the electroporated cells. The mixture was transferred to 1.5 mL microcentrifuge tubes and incubated at 37 °C with shaking for 2 hours to allow recovery. Recovered cells were plated onto Tryptic Soy Agar (TSA) plates containing the appropriate antibiotics. Successful transformation in *S. aureus* was confirmed by colony PCR and agarose gel electrophoresis, using the same conditions previously applied to transformed *E. coli* (**Table 1**).

### Cell culture

THP-1 human monocytic cells (ATCC, TIB-202) were cultured in T25 (10 mL) or T75 (30 mL) tissue culture flasks (Merck) at 37 °C in a humidified incubator with 5% CO₂ and kept up to passge 10. Cells were maintained in RPMI 1640 medium without phenol red (Thermo Fisher Scientific), supplemented with 10% fetal bovine serum (FBS; Thermo Fisher Scientific). Culture medium was freshly prepared prior to each use. Cells were subcultured every 3 days to maintain a density between 1× 10⁵ and 1 × 10⁶ cells/mL. For routine passaging, cell suspensions were transferred into pre-warmed medium to seed new cultures at a density of 1-2 × 10⁵ cells/mL in a total volume of 30 mL in T75 flasks. Cell viability was assessed by trypan blue exclusion (Thermo Fisher Scientific), and viable cells were counted using a TC20 cell counter machine (BioRad).

### Infection of THP-1 with knock-down mutants and staining for microscopy

To investigate the role of *mnhB* in *S. aureus* intracellular persistence, we quantified induced mutant vs. uninduced mutant internalization in THP-1 monocytes using fluorescence microscopy. Knockdown strains were pre-induced for 5 h in RPMI 1640 supplemented with 10% FBS, 10 ng/mL anhydrotetracycline (aTc), and/or 10 µg/mL chloramphenicol. Cultures were grown in 24-well plates (1 mL/well) at 37 °C, 5% CO₂. Uninduced controls (−aTc) were included for each strain. THP-1 monocytes were stained with 2.5 µM CellTracker Green (Thermo Fisher Scientific, cat# C7025) for 20 min at 37 °C and 5% CO_2_, washed, and seeded onto poly-D-lysine–coated 8-well µ-slides (50 µg/mL; P6407, Sigma-Aldrich; 80827, Ibidi). Cells were allowed to adhere for 20 min at 37 °C, 5% CO₂. Bacterial cultures were washed and passed through 3 µm filters (PluriSelect, cat# 46-10003-15) to remove aggregates, then diluted to 2 × 10⁶ CFU in 200 µL in cell culture medium media. Cell culture medium was removed and replaced with bacterial suspension to achieve an MOI of 10. Infections were performed for 10, 30 min or 1 h at 37 °C, 5% CO₂. Post-infection, wells were washed with PBS to remove extracellular bacteria and fixed in fixation buffer (420801, BioLegend) for 20 min at room temperature. Cells were permeabilized and blocked with PBS-T (PBS + 0.1% Twee20) supplemented with 5% goat serum for 1 h in the dark. Infected cells were then stained with 0.5% DAPI and 0.5% Vancomycin–JF669 (Jantarug et al. 2024) for 20 min, followed by three PBS washes.

### Acquisition and analysis of microscopy data

Imaging of infected THP-1 monocytes was performed using a Nikon Crest V3 spinning disk confocal microscope at 40× or 60× magnification. Excitation wavelengths were 405 nm (DAPI), 477 nm (CellTracker), and 638 nm (Vancomycin– JF669). Exposure times were set to 10 ms (brightfield), 250 ms (DAPI), 50 ms (CellTracker), and 1 s (Vancomycin–JF669). To minimize selection bias, five random fields of view were selected per well based on brightfield images alone. Z-stack imaging was performed for each field to capture the full cell volume, starting below and ending above the cells. Z-step intervals were set to 0.5 µm to ensure accurate detection of intracellular *S. aureus*, accounting for the ∼1 µm diameter of the bacterium. Spinning disk confocal acquisition was used for all fluorescence channels to reduce photobleaching and background noise. Images were processed and analyzed using FIJI (ImageJ). Bacteria were defined as intracellular only if their fluorescence (Vancomycin–JF669) co-localized with the cytoplasmic signal (CellTracker). For each condition, the total number of THP-1 cells and the number of infected cells were quantified to calculate infection rates. Additionally, intracellular bacterial volume per monocyte was measured. To improve segmentation, a Gaussian blur filter was applied to reduce background noise. Composite images were split into individual channels (brightfield, DAPI, CellTracker, and Vancomycin–JF669), and background subtraction was applied to the CellTracker and Vancomycin–JF669 channels. Thresholding was used to generate binary masks, enabling segmentation of cells and bacteria. To accurately count individual THP-1 cells and quantify intracellular *S. aureus*, touching or overlapping cells were separated using the “Distance Transform Watershed 3D” function from the MorphoLibJ library in FIJI/ImageJ. This function not only segments adjacent cells but also assigns a unique pixel intensity value (label) to each cell. To exclude incomplete cells at the image borders, the “Kill Borders” function was applied, which removes objects touching the image boundaries in all dimensions, including the Z-axis. The “Connected Components Labeling” function was then used to assign a unique integer label (starting from 1) to each segmented cell, with all pixels belonging to the same object receiving the same value. To identify intracellular bacteria, the Vancomycin-JF669 channel (bacterial signal) was binarized by dividing all pixel values by 255, setting background pixels to 0 and bacterial pixels to 1. The resulting binary mask was then multiplied with the labeled cell image using the “Image Calculator” function. This operation retains only bacteria that overlap with labeled cell regions, effectively removing extracellular bacteria. Intracellular bacteria inherit the label of the host cell, allowing direct linkage between host and pathogen. Quantitative analysis was performed using the “Analyze Regions 3D” function for both the cell and intracellular bacterial images. The output consists of two tables: one listing each cell’s label and volume, and another listing intracellular bacterial clusters and their associated host cell labels. Bacterial clusters sharing the same label are considered to reside within the same host cell, enabling quantification of both infection frequency and bacterial burden per cell. To streamline downstream analysis, a custom R script was used to filter and summarize the data. Cells with a volume above 5,000 voxels (equivalent to 197.8 µmS) and bacterial clusters below 10 voxels (0.524 µmS) were excluded. Bacteria associated with excluded cells were also removed. The script then calculated the number and proportion of infected cells based on the filtered dataset.

### Killing curves

Bacterial inoculum was prepared using a McFarland 0.5 standard. For co-culture experiments, THP-1 cells were infected at a multiplicity of infection (MOI) of 10. Cells were incubated for 1 h at 37 °C in 5% CO₂ to allow internalization. After 1 h, antibiotic was added to the cultures. Colony forming units (CFU) were determined at 0 h (prior to antibiotic addition) and at 2, 4, 6, 8, and 24 h. At each time point, extracellular bacteria were carefully removed, and the cells were washed twice with PBS before being lysed in PBS supplemented with 0.1% Triton X-100 (Sigma). Lysates were plated for CFU enumeration using an Optitron robotic plating system.

## Author contributions

A.I, M.V.M, J.V.W, N.K designed the research. A.I, E.D, M.V.M performed the experiments. R.K provided the clinical information regarding antibiotic concentrations. K.J and P.R.F provided the JF669 probe. A.S.Z provided the *S. aureus* isolates. V.d.B provided feedback on the data analysis and helped with data representation. A.I analysed the data. A.I wrote this manuscript with feedback from all authors. D.B, A.S.Z, J.W.V and N.K supervised this project. N.K, A.S.Z. and J.W.V provided funding.

### Acknowledgements

We thank the microscopy facility of the Department of Biomedicine for their valuable help. We thank all the members from the Khanna lab for their feedback. We thank Alejandro Mejia Gomez for his feedback on the manuscript. V.d.B was supported by an SNSF PostDoc Mobility fellowship (P500PB_225439). This work was funded by the Swiss National Science Foundation (NCCR AntiResist, grant no. 180541 and SNSF project grant 310030_204343 to A.S.Z).

