## Supplemental information for "*Staphylococcus aureus* Mnh1 cation–proton antiporter promotes phagocytic uptake and intracellular survival in human monocytes"

**Supplementary Figures**

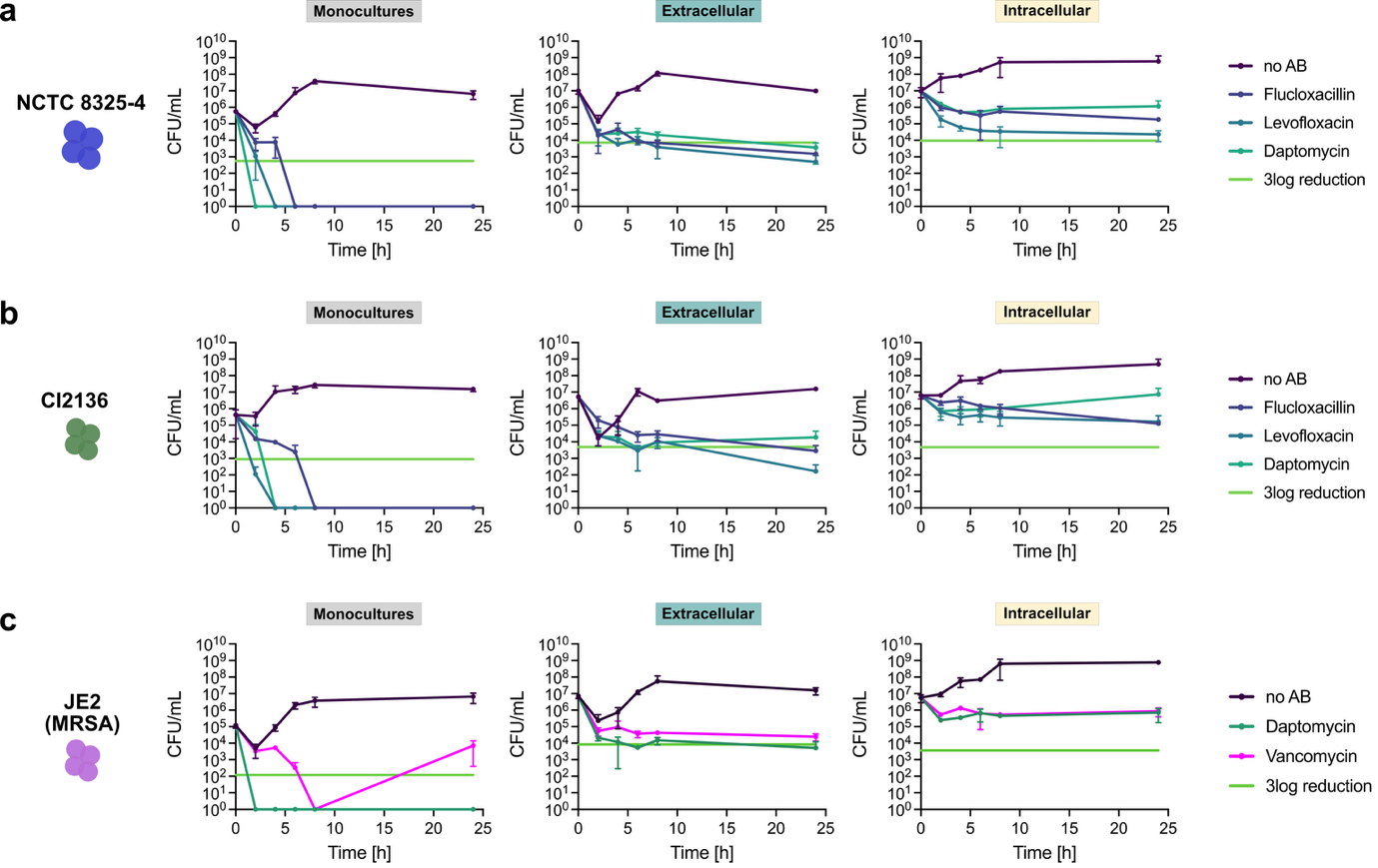

**Figure S1 | Intracellular protection against clinically relevant antibiotics in three S. aureus isolates.** **(a)** NCTC 8325-4, **(b)** CI2136, **(c)** JE2 (MRSA). Growth curves were generated to evaluate S. aureus survival over 24 h in monoculture and within extracellular and intracellular compartments, with or without antibiotic treatment. THP-1 cells were infected at a multiplicity of infection (MOI) of 10 and incubated for 1 h; parallel monocultures were established using the same inoculum. Following infection, cultures were centrifuged, supernatants removed, and fresh medium—with or without antibiotics—was added. Antibiotics were applied at C_max_ concentrations, as defined by the Sanford and/or Mensa guidelines (**Table S1**). For the MRSA strain JE2, which is resistant to levofloxacin, daptomycin and vancomycin were used instead. Colony-forming units (CFUs) were enumerated at 2, 4, 6, 8, and 24 h post-treatment.

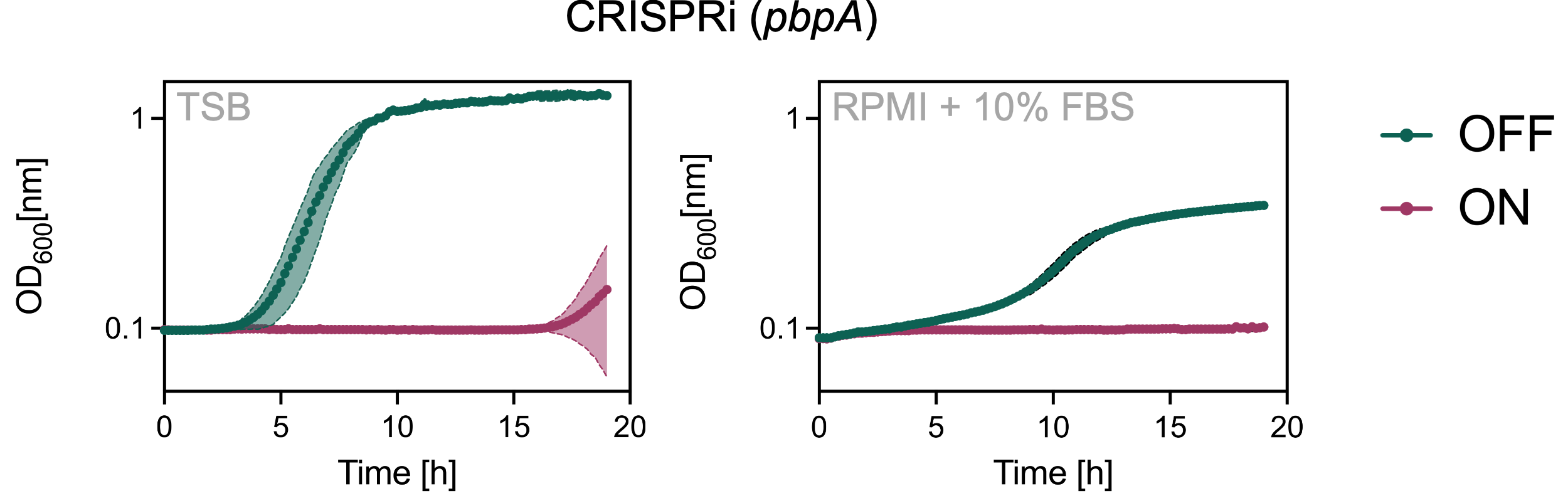

**Figure S2 | CRISPRi system activity in TSB compared with RPMI + 10% FBS.**
Left panel: Growth curves of S. aureus Cowan I *sep*::Ptet-*dcas9* carrying a sgRNA targeting pbpA in TSB, with (CRISPRi ON) or without (CRISPRi OFF) addition of aTc 10 ng/mL. Right panel: Growth curves under the same conditions in RPMI supplemented with 10% FBS, showing a reduced growth rate compared to TSB. Shaded areas represent ± standard deviation across n = 3 replicates.

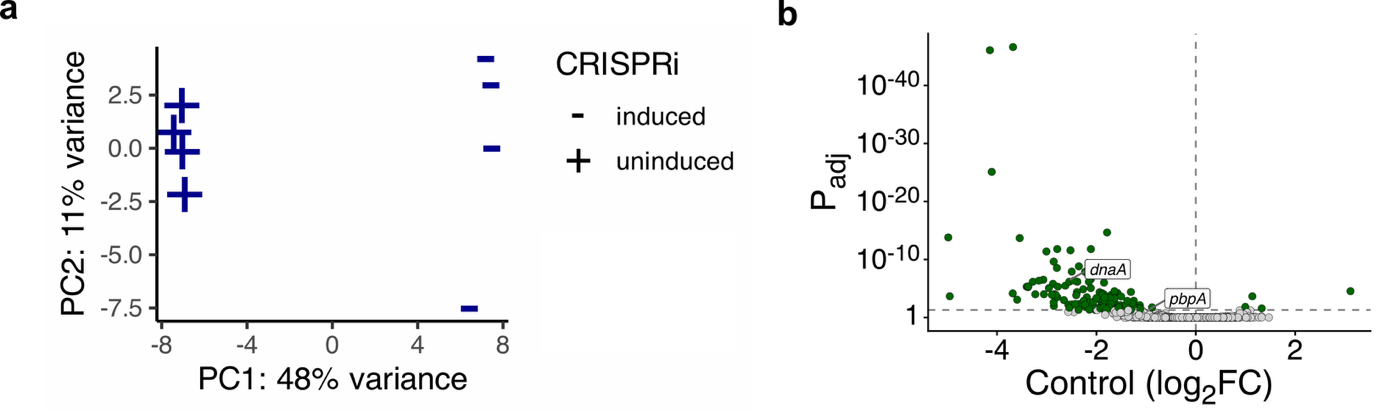

**Figure S3 |** Cowan I CRISPRi library grown in RPMI supplemented with 10% FBS, without THP-1 cells, used as a control. **(a)** Principal component analysis (PCA) illustrating clear separation between induced (+aTc, 10 ng/mL) and uninduced control samples (no aTc). **(b)** Volcano plot displaying the differential analysis between uninduced and aTc-induced libraries in the control condition, with *dnaA* highlighted as a representative essential gene.

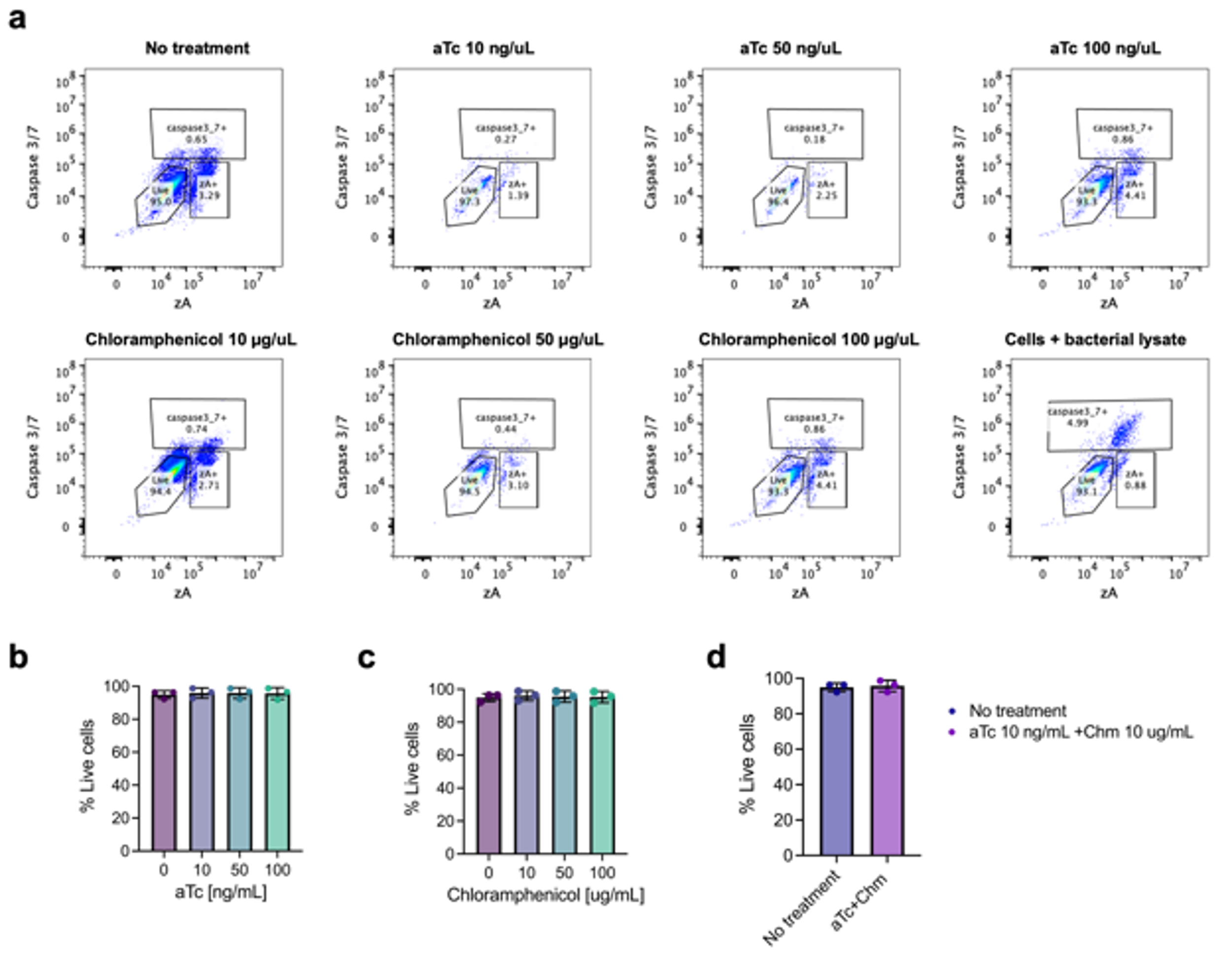

**Figure S4 | CRISPRi screen growth conditions did not induce toxicity in THP-1 cells.**
THP-1 cells were cultured in RPMI supplemented with 10% FBS and either anhydrotetracycline (aTc) or chloramphenicol for 6 h (1 h longer than the screen performed). Cell viability was assessed by flow cytometry using Zombie Aqua (zA) staining, and apoptosis was evaluated using a caspase-3/7 marker. **(a)** Representative gating strategy for each sample. **(b)** Percentage of live cells at different aTc concentrations. **(c)** Percentage of live cells at different chloramphenicol concentrations. **(d)** Percentage of live cells under the co-culture condition used in the CRISPRi screen (10 ng/mL aTc and 10 µg/mL chloramphenicol). Data represent three independent experiments.

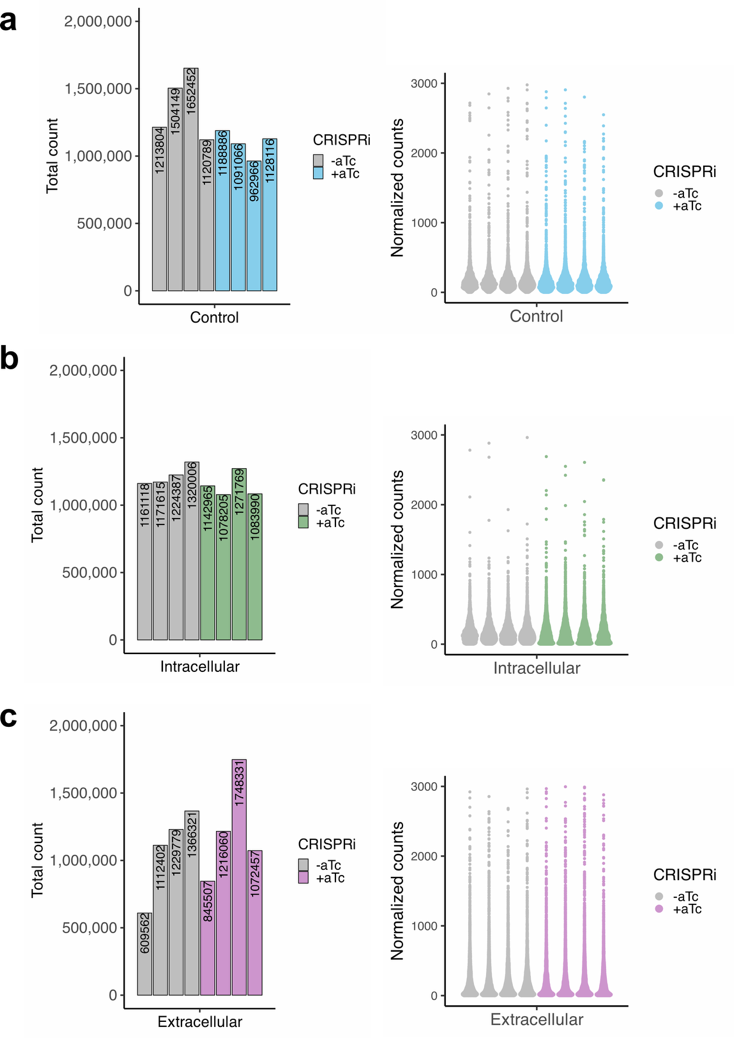

**Figure S5 |** **(a)**  sgRNA count totals and normalized sgRNA count distributions for the control condition (RPMI + 10% FBS without cells), with and without dCas9 induction using anhydrotetracycline (aTc). **(b)** Raw and normalized sgRNA count distributions for the intracellular condition, with and without dCas9 induction using anhydrotetracycline (aTc). **(c)** Raw and normalized sgRNA count distributions for the extracellular condition, with and without dCas9 induction.

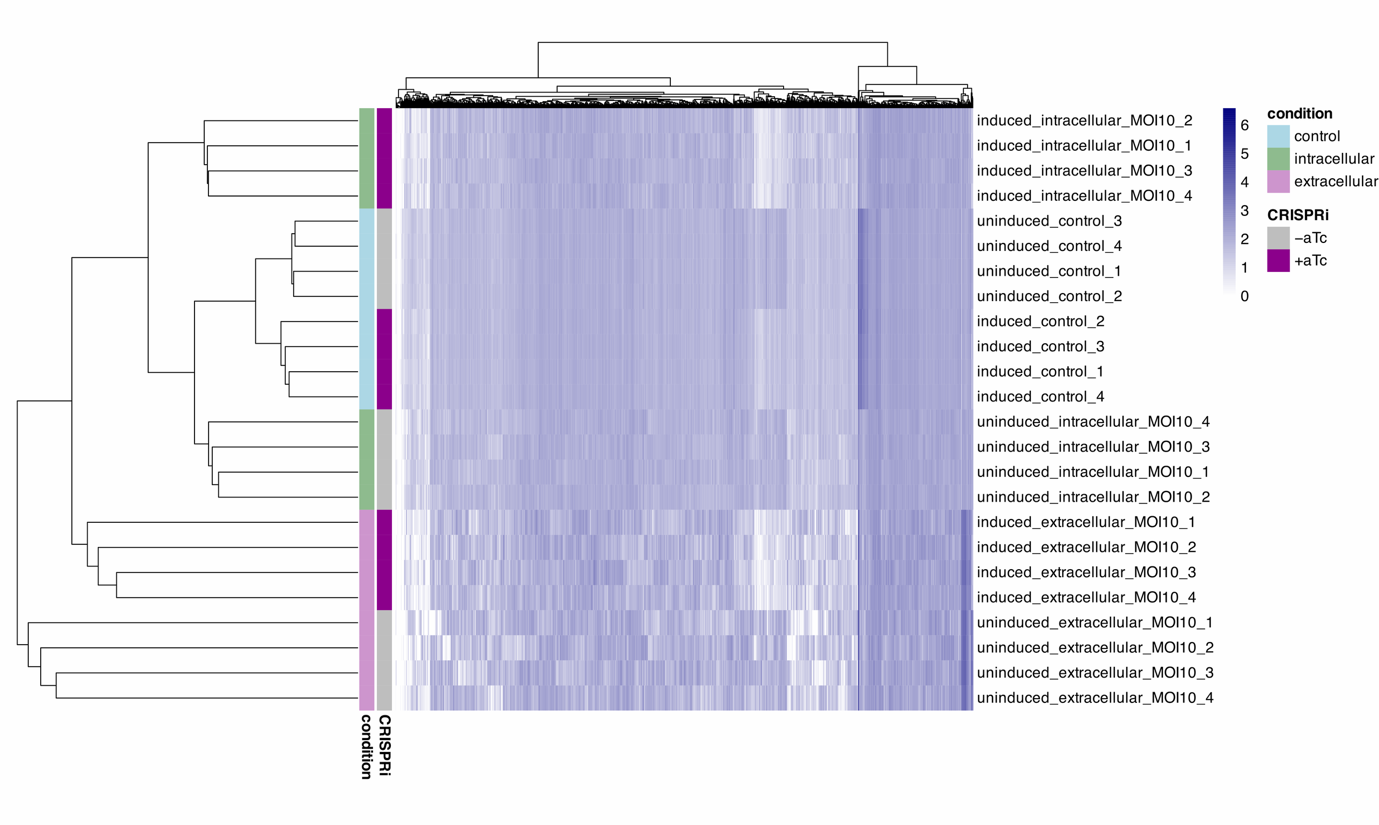

**Figure S6 |** Heatmap of sgRNA enrichment across all technical replicates for control, intracellular, and extracellular conditions under induced and uninduced treatments. Data are shown as log₁₀(normalized count + 1) to avoid undefined values at zero and to enhance visualization of low-abundance (essential) sgRNAs.

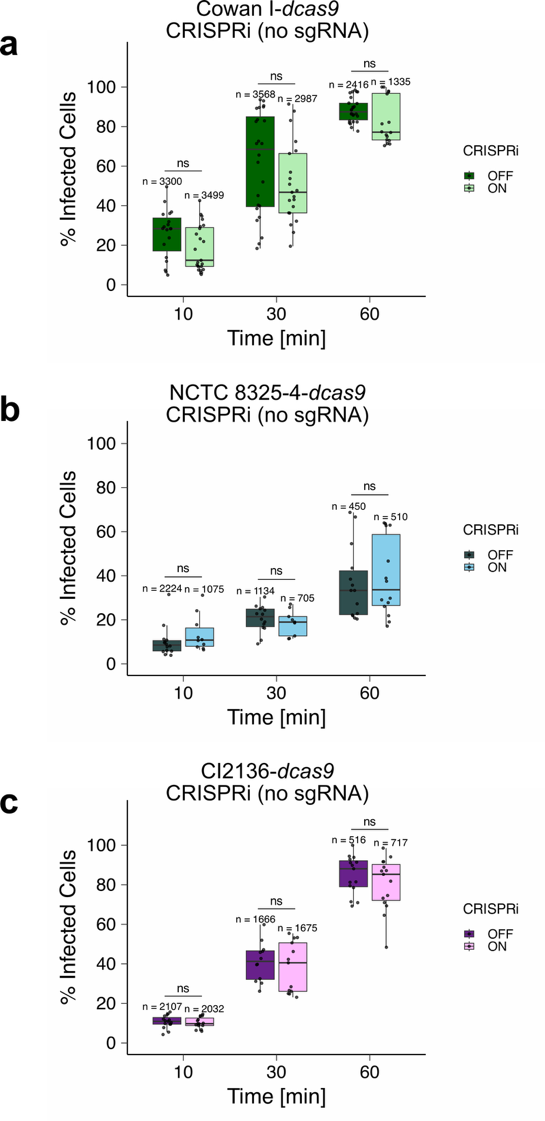

**Figure S7 | Control experiment assessing the effect of aTc on internalization percentage.**

(a)  Cowan I, (b) NCTC 8325-4, and (c) clinical isolate CI2136. Each strain carries an intrachromosomal ***dcas9*** but no sgRNA, serving as a control for the effect of aTc on internalization. Each data point represents the mean percentage of infected cells per field of view, each fields containing ≥70 cells. For each condition, 3–5 images were acquired per experiment, and experiments were independently repeated 3–4 times. Statistical significance was determined using the Wilcoxon rank-sum test. Asterisks denote p-values: *** < 0.001, ** < 0.01, * < 0.05; ns, not significant. The total number of cells quantified per condition is shown above each boxplot.

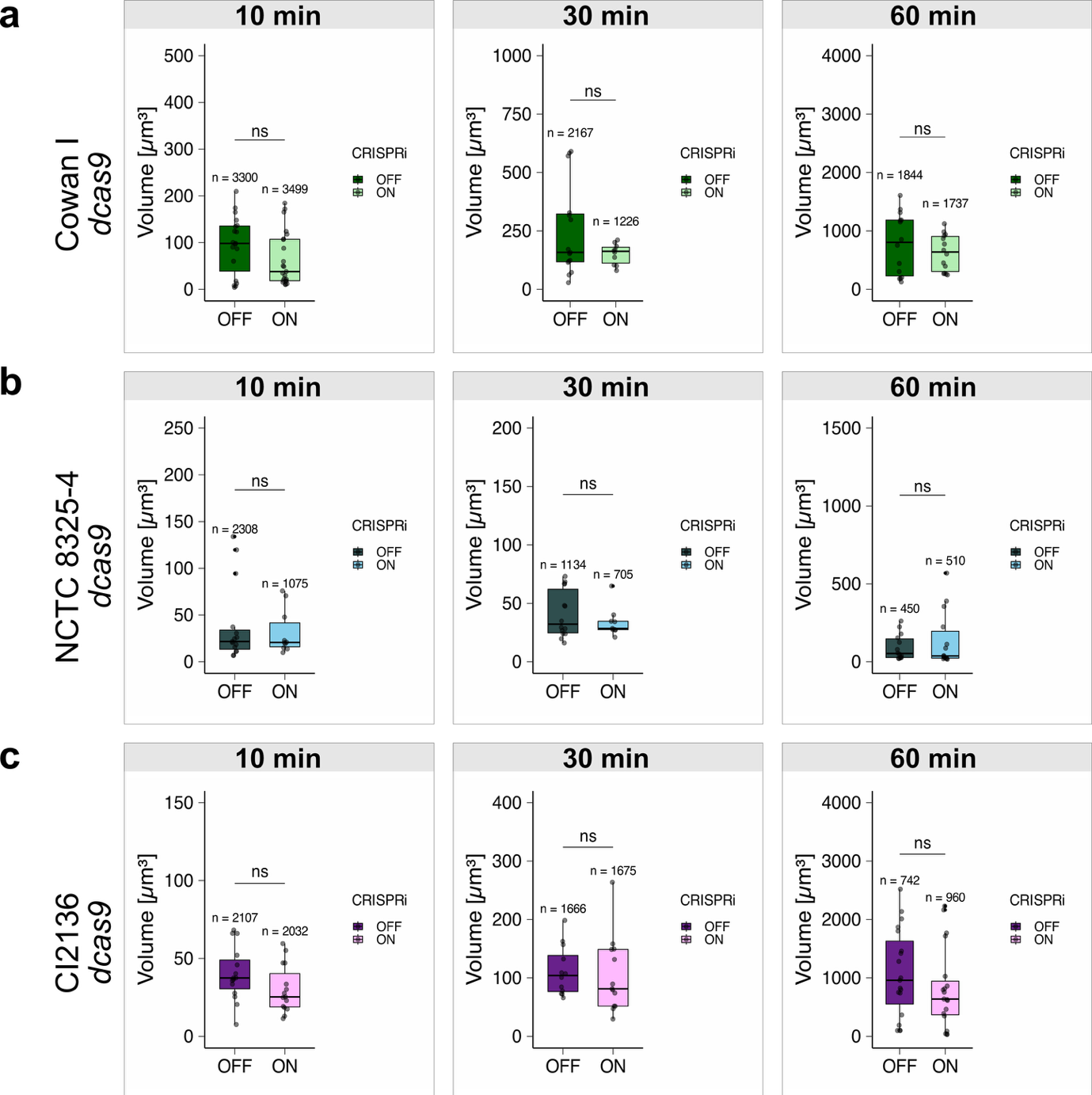

**Figure S8 | Control experiment assessing the effect of aTc on intracellular bacterial volume.** (a)  Cowan I, (b) NCTC 8325-4, and (c) clinical isolate CI2136. Each strain carries an intrachromosomal ***dcas9*** but no sgRNA, serving as a control for the effect of aTc on the volume of bacteria internalized by THP-1 cells. Each data point represents the mean percentage of bacterial volume per cell per field of view (≥70 cells/field). For each condition, 3–5 images were acquired per experiment, and experiments were independently repeated 3–4 times. Statistical significance was determined by Wilcoxon rank-sum test. Asterisks denote p-values: *** < 0.001, ** < 0.01, * < 0.05; ns, not significant. The total number of cells quantified per condition is indicated above each boxplot.

**Supplementary tables**

**Table S1|** C_max_ concentrations as reported in **(a)** the Sanford guide^50^ and **(b)** the Mensa guide^49^. **(c)** C_max_ concentrations used in this study, adjusted for protein binding percentages listed in the respective guides. These values were further refined based on antibiotic studies previously conducted in our laboratory (Zurbrügg et al., in prep.; Morin et al., in prep.).

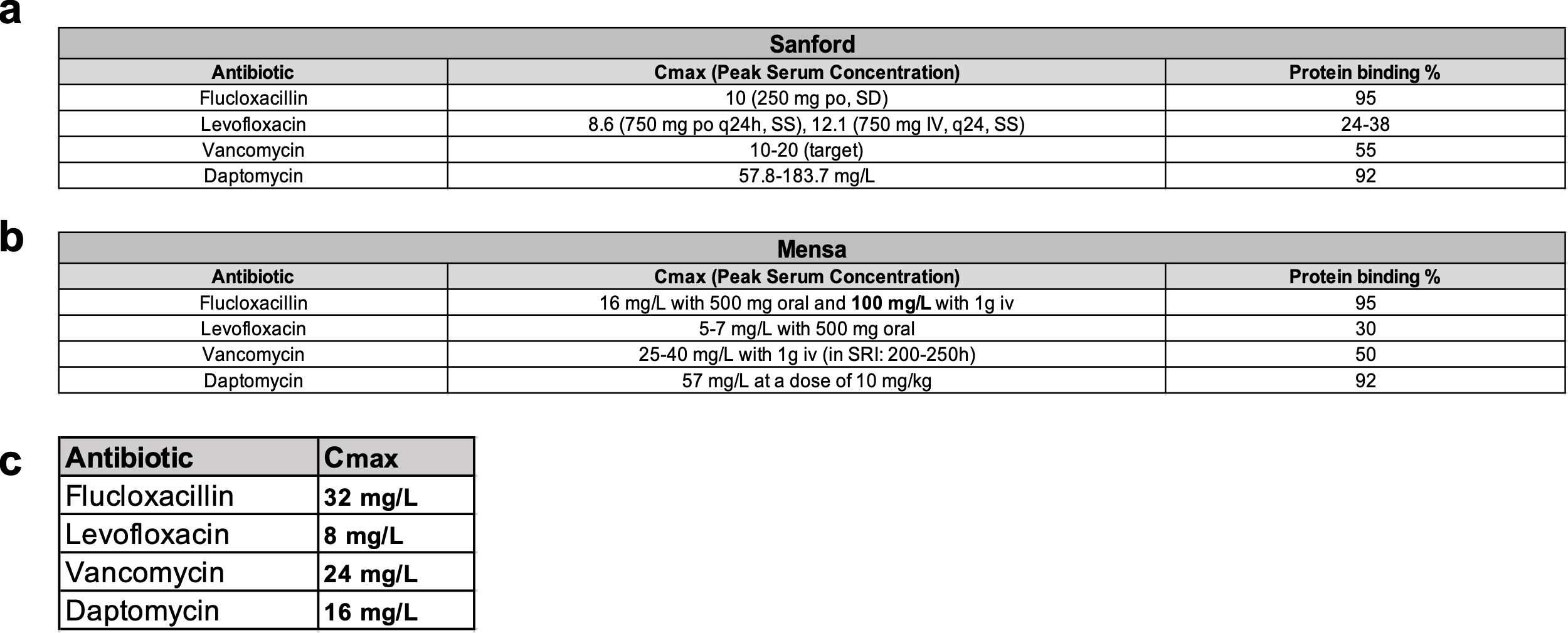

**Table S2|** List of knockdown strains generated in the study. Oligos used for each mutant are indicated.

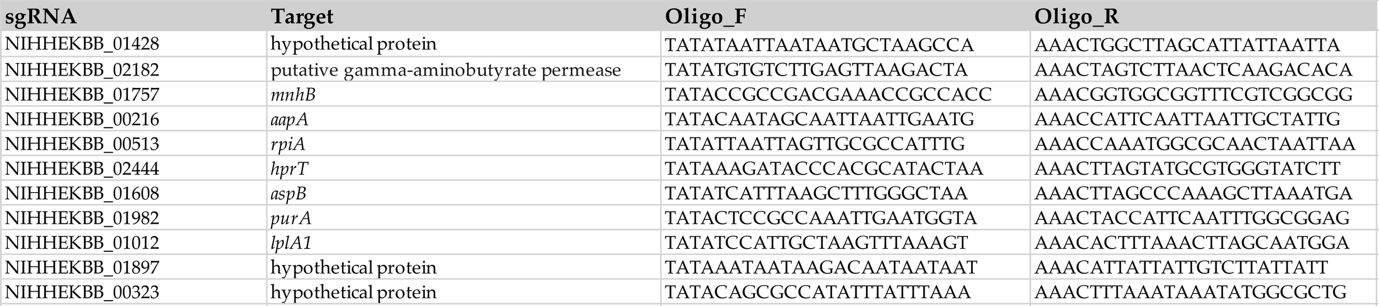

**Table S3|** List of oligos used for cloning sgRNA to generate knockdown strains.

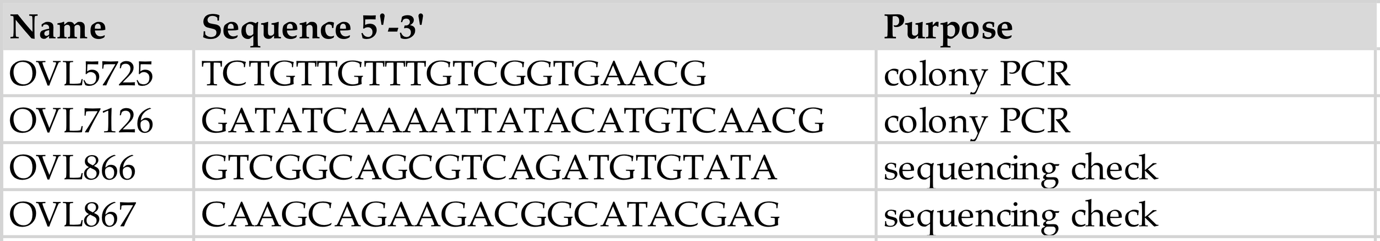
